# The Landscape of Mutational Mosaicism in Autistic and Normal Human Cerebral Cortex

**DOI:** 10.1101/2020.02.11.944413

**Authors:** Rachel E. Rodin, Yanmei Dou, Minseok Kwon, Maxwell A. Sherman, Alissa M. D’Gama, Ryan N. Doan, Lariza M. Rento, Kelly M. Girskis, Craig L. Bohrson, Sonia N. Kim, Lovelace J. Luquette, Doga C. Gulhan, Brain Somatic Mosaicism Network, Peter J. Park, Christopher A. Walsh

## Abstract

Although somatic mutations have well-established roles in cancer and certain focal epilepsies, the extent to which mutational mosaicism shapes the developing human brain is poorly understood. Here we characterize the landscape of somatic mutations in the human brain using ultra-deep (~250×) whole-genome sequencing of brains from 59 autism spectrum disorder (ASD) cases and 15 controls. We observe a mean of 26 (±10, range 10-60) somatic single nucleotide variants (sSNVs) per brain present in ≥4% of cells, with enrichment of mutations in coding and putative regulatory regions. Our analysis reveals that the first cell division after fertilization produces ~3.4 mutations, followed by 2-3 mutations in subsequent generations. This rate suggests that a typical individual possesses ~80 sSNVs present in ≥2% of cells—comparable to the number of *de novo* germline mutations per generation—with about half of individuals having at least one potentially function-altering somatic mutation somewhere in the cortex. Although limited by sample size, ASD brains show an excess of somatic mutations in neural enhancer sequences compared to controls, suggesting that mosaic enhancer mutations may contribute to ASD risk.

## Introduction

Somatic mutations, also referred to as mosaic mutations, are acquired post-fertilization and are present in a subset of an individual’s cells, marking only cells that are descended from the originally mutated cell^1^. Mutations occurring early in development may be present throughout the body, whereas later-occurring mutations are found in progressively smaller subsets of cells. Somatic mutations can be neutral or they can confer a functional advantage or disadvantage to the affected cell. Somatic mutations represent a particularly interesting phenomenon in the brain, as most neurons are postmitotic and will harbor their mutations for the life of the individual. The study of benign clonal somatic mutations in the brain has the potential to reveal insights into mechanisms of normal neurodevelopment. Although function-altering somatic mutations have well-established roles in cancer and some focal epilepsies^2–4^, the question of whether mutational mosaicism affects risk of other neuropsychiatric diseases is only beginning to be explored^5–7^.

Previous studies have established that single human neurons harbor large numbers of somatic mutations^8^. However, the burden and extent of *clonal* somatic mutations—somatic mutations present in multiple cells and thus likely to have capacity for functional impact—have until now been difficult to characterize. This is largely due to the fact that most studies of clonal somatic mutation in human brain have been performed in very small sample sizes, based on exomes only^9,10^, or with insufficient depth of sequencing. Investigating the genome-wide landscape of clonal somatic mutation within the human brain is crucial in order to better understand the ways in which developmental mutations shape the adult brain, contribute to normal variation, and cause disease.

ASD is a complex and heterogeneous neurodevelopmental disease characterized by impairments in communication and social interactions as well as repetitive behaviors. Several large studies of blood or saliva DNA from ASD families have shown that both *de novo* germline mutations and exonic sSNVs play a role in ASD causation, with anywhere from 5.4-22% of mutations that were previously thought to be *de novo* actually representing postzygotic mutations^11–14^. Due to the scarcity of donated brain tissue, very few studies have investigated somatic mutation in autism brain^12,15^. Until now, deep whole-genome sequencing (WGS) of autism brain has never been published, and therefore the critical question of how non-coding somatic mutations in autism brain might impact disease risk remains unanswered.

Here we present ultra-deep WGS of 59 ASD brains and 15 neurotypical brains, representing the largest such collection ever assembled. A prior study of ASD brain examined targeted sequencing of 78 genes^15^, whereas we perform whole-genome analysis of prefrontal cortex from some of the same brains and many additional ones. Our deep WGS data enable detection of sSNVs occurring early in development— mutations that are most likely to alter brain function compared to later-occurring mutations—and accurate estimation of allele fractions, allowing for the highest-resolution analysis of human brain mutational mosaicism to date. Our analysis provides unprecedented insight into the general architecture and mutational signatures of somatic mutation in the normal human brain, as well as important implications for genome-wide mosaic mutation in ASD pathogenesis.

## Results

### Variant discovery and validation

We sampled 59 ASD brains and 15 controls (Supplementary Table 1), extracting DNA from dorsolateral prefrontal cortex (PFC) where available, before performing WGS to an average depth of 250×. The majority of ASD samples and all control samples were sequenced using a PCR-free library preparation, whereas 11 ASD samples were sequenced with a PCR-based preparation. We identified sSNVs and indels in all samples using a new machine learning-based method called MosaicForecast^16^, which is optimized to detect somatic mutations in the absence of a matched reference tissue (Fig. 1, Supplementary Fig. 1 and 2). This is crucial for studying disease samples such as ASD brains, as many donated cases do not include paired non-brain tissues. We were able to call mutations with variant allele fraction (VAF) down to 2% in PCR-free samples, and to 3% VAF in PCR-based samples; therefore, we had sensitivity to detect variants present in as few as 4-6% of cells in a given section of brain tissue. After stringent filtration, we identified an average of 25.6 ± 10.2 sSNVs per sample (range 10 to 60, Fig. 2a, Supplementary Table 2). VAFs of detected sSNVs ranged from ~2-39%, corresponding to mutation presence in ~4-78% (mean 17%) of cells for autosomal variants (Fig. 2b). Identified sSNV positions were covered at an average of 216X.

**Figure 1:**
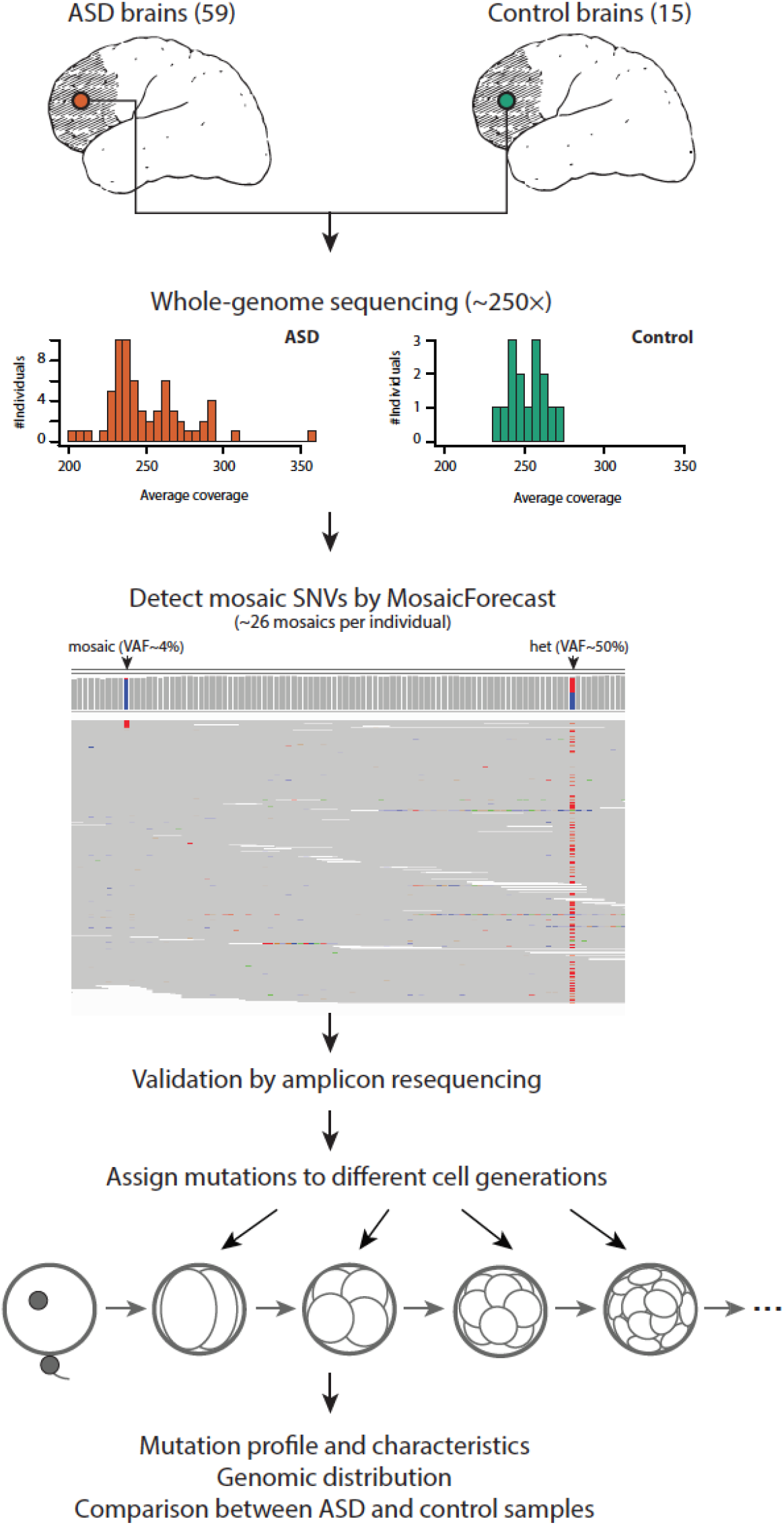
Experimental design and genome coverage. DNA was isolated from dorsolateral prefrontal cortex of 59 ASD brains and 15 control brains then whole-genome sequenced to an average depth of 250×. Germline variants were identified and somatic single nucleotide variants were called using MosaicForecast. A representative set of mutations was validated using targeted amplicon resequencing. Mutations were systematically assigned to cell generations based on variant allele fraction.

Deep targeted resequencing of 208 putative sSNVs (Supplementary Table 3), to an average depth of ~50,000× per reaction, demonstrated the accuracy of our mutation-calling algorithm since called sSNVs showed an overall validation rate of 90% (Fig. 2c; 94% for samples sequenced using PCR-free libraries). VAFs from targeted resequencing were highly concordant with VAFs estimated from WGS (Fig. 2d, R^2^=0.904). Analysis of germline heterozygous sites covered in deep resequencing showed overdispersion of validation VAFs, which was not present in WGS data and was corrected in downstream analyses (Supplementary Fig. 3, Supplementary Table 4). Since the 11 samples sequenced using PCR-based library preparation had a lower validation rate (Fig. 2c), only sSNVs discovered in PCR-free samples or validated in PCR-based samples were used for downstream analyses. We also validated 86 mosaic indels called with MosaicForecast (Supplementary Table 5, Supplementary Fig. 4), which were used in regulatory region analyses as described later.

**Figure 2:**
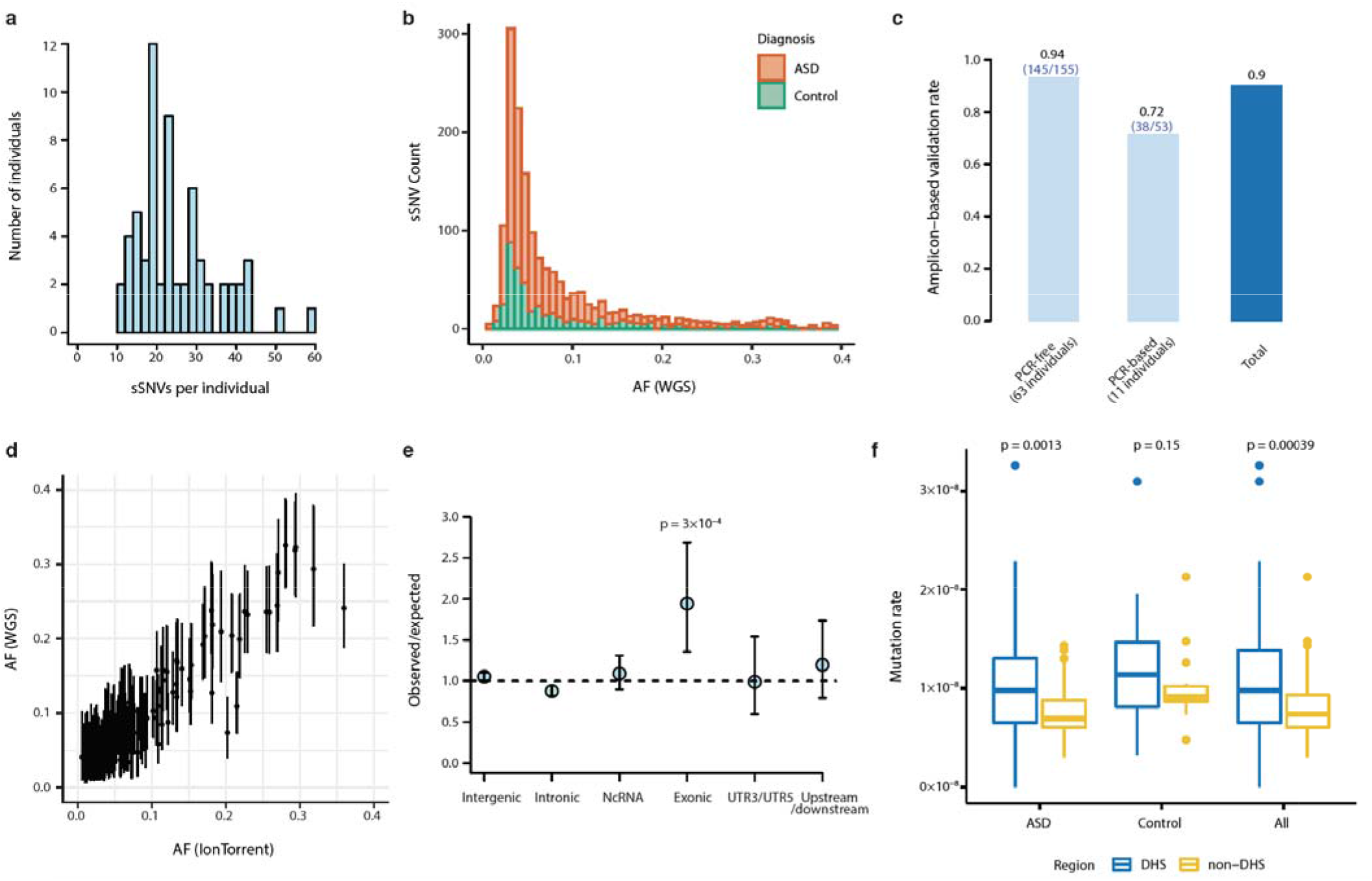
Mosaic mutations are present across the genomes of cases and controls. **a**, Distribution of mosaic mutations per subject. **b**, VAF distribution of all mosaic variants identified in this study (stacked barplot). There are more ASD cases than controls and therefore more total ASD sSNVs, but no difference in allele fraction distribution. **c**, Validation rates in PCR-free and PCR-based samples from deep targeted resequencing of 208 putative mutations. **d,** Variant allele fractions from WGS were highly correlated with allele fractions from deep resequencing (R^2^=0.904). **e,** Number of observed mosaic mutations divided by the expected number of mosaics assuming a uniform mutation rate. Error bars indicate 95% CIs calculated with a binomial test. Exonic regions show the strongest enrichment for mosaics among several genomic regions. **f**, Non-coding somatic mutations are enriched in DNAse I hypersensitive sites annotated by the Roadmap Epigenomics Project in both ASD cases and the total dataset. The lower and upper hinges of boxplots in panels d, e, and f correspond to the first and third quartiles, and the middle lines correspond to the median values.

### Somatic mutations are enriched in coding and open chromatin regions

At the single-neuron level, it has been shown that sSNVs are enriched in exons, suggesting that transcriptional error may contribute to somatic mutation^17^. However, few studies have examined clonal somatic mutations genome-wide in bulk DNA samples to determine which regions of the genome harbor somatic mutations that arise in early development. We found that across all PCR-free samples, 35 sSNVs were exonic (2.2%), which is about twice as high as expected (Fig. 2e, p=0.0003, two-tailed binomial test, Supplementary Table 6). Among these, 12 were silent, 2 were protein-truncating, and 21 were missense. Approximately 43% (27/63) of PCR-free samples had at least one detectable exonic sSNV, with one sample having three exonic mutations (Supplementary Fig. 5) at VAFs detectable in this study. These data suggest that coding regions are particularly vulnerable to somatic mutation during development.

Across all samples, we identified 21 potentially damaging exonic sSNVs, including validated variants in PCR-based samples (Table 1, Supplementary Table 6). Damaging somatic mutations in mutationally constrained genes (as predicted by pLI score) were identified in both cases and controls based on rigorous criteria using 12 different mutation effect prediction tools^18^, with the classification of NsynD4 representing the most likely damaging nonsynonymous mutations and the classification of LOF representing predicted loss-of-function mutations (Table 1). Our dataset included two protein-truncating mutations predicted with high confidence to be loss-of-function, though both were in genes predicted to be relatively tolerant to loss-of-function (Supplementary Table 6).

**Table 1:** Potentially damaging exonic sSNVs identified in this study.

| Gene | Mutation (cDNA) | Mutation (Protein) | Case ID | Diagnosis | Library Prep Method | VAF | Validation | Pathogenicity Prediction | pLI Score | Missense Z score | SFARI Gene Score | Other Disease Association |
| --- | --- | --- | --- | --- | --- | --- | --- | --- | --- | --- | --- | --- |
| **Damaging Mosaic Mutations in Constrained Genes** |  |  |  |  |  |  |  |  |  |  |  |  |
| CACNA1A | NM_001127221.1:c.118G>T | p.G40W | UMB1174 | ASD | PCR-Free | 0.055 | Mosaic | NsynD4 | 1 | 7.23 | S (syndromic evidence) | Autosomal dominant epileptic encephalopathy |
| SAFB2 | NM_014649.2:c.2120G>A | p.R707H | UMB1174 | ASD | PCR-Free | 0.058 | Mosaic | NsynD4 | 0.99 | 1.32 | - |  |
| PTPN12 | NM_001131009.1:c.437C>T | p.A146V | MSSM001 | ASD | PCR-Free | 0.042 | Mosaic | NsynD4 | 1 | -0.05 | - | Somatic colon cancer |
| DCAF8 | NM_015726.3:c.1585C>T | p.R529W | UMB4899 | ASD | PCR-Based | 0.039 | Mosaic | NsynD4 | 1 | 3.73 | - | Autosomal dominant giant axonal neuropathy |
| TCERG1 | NM_001040006.1:c.1829A>G | p.D610G | UMB4334 | ASD | PCR-Based | 0.289 | Mosaic | NsynD4 | 1 | 3.62 | - |  |
| COL11A2 | NM_080679.2:c.4367C>T | p.T1456I | UMB4899 | ASD | PCR-Based | 0.044 | Mosaic | NsynD4 | 1 | 2.04 | - | Deafness |
| AGAP3 | NM_001308305.1:c.818C>G | p.P273R | UMB5841 | ASD | PCR-Free | 0.046 | N/A | NsynD4 | 0.99 | 4.35 | - |  |
| FAM13B | NM_001101800.2:c.1682A>G | p.D561G | UMB5176 | ASD | PCR-Based | 0.047 | Mosaic | NsynD4 | 0.9 | -0.53 | - |  |
| EIF4G3 | NM_003760.4:c.4634C>T | p.A1545V | UMB4842 | Control | PCR-Free | 0.044 | Mosaic | NsynD4 | 1 | 2.3 | - |  |
| AEBP2 | NM_001114176.1:c.1057T>A | p.F353I | UMB1712 | Control | PCR-Free | 0.047 | Mosaic | NsynD4 | 0.95 | 3.74 | - |  |
| **Damaging Mosaic Mutations in Non-Constrained Genes** |  |  |  |  |  |  |  |  |  |  |  |  |
| DNAH3 | NM_001347886.1:c.8578G>A | p.E2860K | AN09412 | ASD | PCR-Free | 0.127 | Mosaic | NsynD1 | 0 | -1.74 | 4 (minimal evidence) |  |
| SCARF1 | NM_003693.3:c.1943C>T | p.P648L | AN09412 | ASD | PCR-Free | 0.058 | Mosaic | NsynD2 | 0 | 0.63 | - |  |
| CAST | NM_001042440.4:c.2T>G | p.M1R | M3663M | ASD | PCR-Free | 0.04 | N/A | NsynD2 | 0 | -1.8 | - | Autosomal recessive dermatologic disease |
| COL6A3 | NM_057166.4:c.988G>A | p.E330K | UK25363 | ASD | PCR-Free | 0.031 | Mosaic | NsynD4 | 0 | -0.3 | - |  |
| PARVA | NM_018222.4:c.1184A>G | p.K395R | UMB1174 | ASD | PCR-Free | 0.02 | N/A | NsynD4 | 0.34 | 0.98 | - |  |
| GLT8D2 | NM_001316967.1:c.809C>T | p.T270I | UMB4849 | ASD | PCR-Free | 0.04 | Mosaic | NsynD4 | 0 | -0.29 | - |  |
| FAM83E | NM_017708.3:c.1336C>T | p.R446X | UMB914 | Control | PCR-Free | 0.057 | Mosaic | LOF-1* | 0 | -0.38 | - |  |
| UNC45A | NM_001323620.1:c.1416G>C | p.K472N | UMB1024 | Control | PCR-Free | 0.062 | Mosaic | NsynD4 | 0 | -0.38 | - |  |
| RNF175 | NM_173662.2:c.305G>A | p.W102X | UMB1465 | Control | PCR-Free | 0.072 | Mosaic | LOF-1* | 0 | -1.07 | - |  |
| TTN | NM_133378.4:c.17758C>T | p.P5920S | UMB4842 | Control | PCR-Free | 0.158 | Mosaic | NsynD4 | 0 | -5.48 | 4S (minimal and syndromic evidence) | Cardiomyopathy, muscular dystrophy |
| PTGDR | NM_000953.2:c.206C>T | p.T69M | UMB5391 | Control | PCR-Free | 0.047 | Mosaic | NsynD2 | 0 | 1.03 | - |  |

\*Also predicted high-confidence LOF by LOFTEE

Not surprisingly given our small sample size, the overall burden of exonic sSNVs was similar in our ASD cases and controls, but several of our ASD cases carried damaging mosaic mutations that may be relevant for the patient phenotype. For instance, a likely damaging missense mutation in *CACNA1A* (c.354G>T; p.G40W), a gene previously documented to cause autism and intellectual disability in the heterozygous state^19^, was found in approximately 10% of brain cells for case UMB1174 (Table 1, Supplementary Fig. 6). Importantly, *CACNA1A* mutations can also cause epileptic encephalopathy^20^, and case UMB1174 was noted to have a seizure disorder in addition to a diagnosis of ASD. While the study size is not well powered to analyze germline ASD mutations, several individual cases nonetheless showed rare, predicted damaging germline variants in autism risk genes that are likely to contribute to disease, and which may be useful in guiding studies of these samples (Supplementary Table 7).

In addition to finding enrichment of sSNVs in exons, we also observed an excess of somatic mutation in areas of open chromatin. We found an increased rate of somatic mutations in non-coding DNase I hypersensitive sites (DHSs) (p=0.00039, two-tailed Wilcoxon rank sum test, Fig. 2f), which often co-localize with regulatory elements such as promoters and enhancers^21,22^. The increased rate of somatic mutation in DHSs could represent heightened vulnerability of regulatory regions to DNA damage and replication errors during early development^23^.

### Clonal mutation analysis reveals insights into early embryonic development

We examined mutational dynamics in the early embryo by assigning sSNVs to specific cell generations based on their allele fractions (accounting for variable read depth), using a maximum likelihood approach (see Methods, Fig. 3a, Supplementary Table 8). We confirmed that our assigned cell generations correlated well with data from three brains that had previously undergone single-cell lineage analysis^8,17^ (Fig. 3b). When we estimated mutation rate per cell generation after correcting for detection sensitivity, our data revealed an elevated mutation rate during the first post-conception cell division (~3.4 mutations/division), followed by a steady rate of ~2-3 mutations/division in subsequent cell divisions (Fig. 3c; p=0.022 for the difference between the first two cycles based on permutation test, see Methods and Supplementary Fig. 7). We note that our validation rate was high across a wide range of VAFs (validation rate = 100% for mosaics with >0.2 VAF, Supplementary Fig. 8), and analysis of single-cell sequencing data^8^ confirmed that even high-VAF (>0.2) variants were indeed mosaic and not germline (Supplementary Fig. 9). These findings suggest that the higher mutation rate during the first cell generation is unlikely to be artifactual, and it has been suggested before using similar methods^24^.

**Figure 3:**
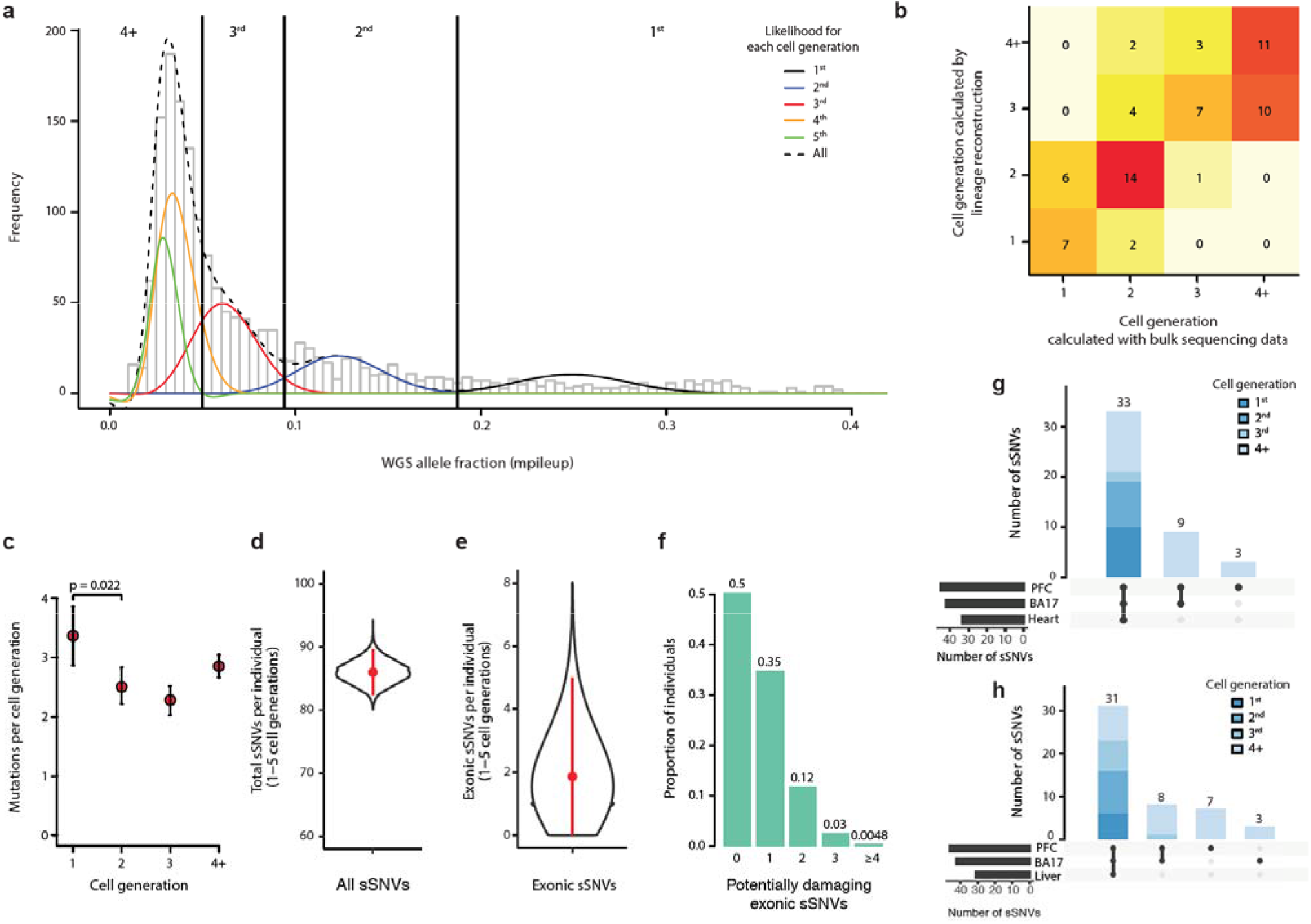
Clonal mutation analysis reveals mutational dynamics in the early embryo. **a**, Clonal somatic mutations map onto a symmetrical model of cell division in the early embryo. The black curve represents the likelihood of sSNVs belonging to first cell generation, blue the second cell generation, red the third cell generation, yellow the fourth cell generation, and green the fifth cell generation. **b**, Cell generation assignments for sSNVs were congruent with data from single-cell lineage analysis of three individuals (UMB1465, UMB4643, UMB4638). **c**, Somatic mutations are elevated in the first cell generation of embryogenesis (~3.4 mutations per cell division), then occur at a rate of approximately 2-3 mutations per cell division in subsequent generations. **d**, Based on the mutation rate per cell generation, an average individual would carry ~86 (95% CI: 82-90) sSNVs from the first five cell generations. **e**, Given that ~2.2% of sSNVs in our dataset are exonic, each individual would carry ~1.9 (95% CI: 0-5) exonic sSNVs. **f**, Assuming ~37% of new exonic mutations are damaging, ~50% of individuals would carry ≥1 damaging exonic mutation from the first 5 cell divisions, present in roughly ≥2% of cells. **g-h**, Amongst mosaic mutations in the occipital lobe and prefrontal cortex for control brains UMB4638 and UMB4643, variants assigned to earlier cell divisions are present in wider tissue distributions across the body.

Based on the positions of the peaks in our mutational VAF distribution, it is possible to infer whether the VAF distribution is more consistent with a symmetric or an asymmetric model of early embryo development. We found that our data slightly favor an asymmetrical cell model (p=3*10^−4^, likelihood ratio test), in which many progenitor cells contribute unevenly to the organism^25^ (Supplementary Fig. 10-11). Importantly, cell generation assignments changed very little with implementation of the symmetrical versus asymmetrical models (Supplementary Table 8).

The estimated early embryo mutation rate is largely consistent with other reports, which have estimated roughly 1-3 mutations per cell division in the early embryo^24–26^. Notably, this somatic mutation rate is about ~1-5 times higher than the germline mutation rate inferred from *de novo* germline mutations in maternal haploids, and ~10 times higher than the estimated germline mutation rate in paternal haploids^24,27^. The cell cycle in the early human embryo is relatively short^28^, with reduced G1 checkpoint protein expression^29^, suggesting one mechanism for the elevated rates of somatic mutation. Furthermore, some mutations in the first cell cycle may represent sites of single-stranded DNA damage in sperm or egg^30,31^, repaired to the mutated base during the first cell division, thereby accounting for the elevated mutation rate in the first cell generation after fertilization.

### ~50% of individuals possess potentially damaging exonic sSNVs in >2% of cortical cells

Every study of somatic mutation is inherently limited by tissue sampling, as somatic mutations by nature are present in some regions of particular organs but not in others. Therefore, it has traditionally been difficult to estimate body-wide or even organ-wide somatic mutation burden. We utilized our cell generation assignments to estimate global somatic mutation burden in the brain.

Assuming early embryo mutation rates as estimated above, a stable mutation rate in the 4^th^ and the 5^th^ cell generations, and no proliferative advantage of variants, a typical individual would amass ~86 (95% CI: 82-90) genome-wide sSNVs in the 1^st^-5^th^ cell generations after conception, each present at ≥1% VAF across the adult body (Fig. 3d). Our directly measured ~26 early embryonic sSNVs, and this estimate of 86 mutations over the first 5 cell divisions, are of quite similar magnitude to the 60 *de novo* germline mutations^44^ per individual identified in a recent WGS family-based study, creating the possibility that function-altering mosaic mutations may contribute to variation more commonly than appreciated.

To estimate this contribution to functional variation we focused on the ~2.2% of observed variants in our dataset that are exonic, suggesting that a typical individual would have on average ~1.9 (95% CI: 0-5) exonic sSNVs (Fig. 3e) occurring in the first five cell divisions of development. Large studies have found that approximately ~45% of new exonic variants that have not yet undergone evolutionary selection are likely to be damaging^32,33^ and among our set of 35 PCR-free exonic mutations, 37% are predicted to be most damaging with scores of either NsynD4 or LOF (Supplementary Table 6). Therefore, assuming ~37% of exonic sSNVs are damaging, then ~50% of individuals would carry ≥1 damaging exonic sSNV at ≥1% VAF (≥2% of cells) (Fig. 3f). Importantly, even low-VAF mutations have potential to cause disease, as damaging mutations with allele fractions as low as 1% and present in only a small area of the brain have been frequently reported to cause epilepsy via focal cortical dysplasia^3,4^. When we include non-coding sSNVs (as we will describe), the number of potentially damaging sSNVs increases. This analysis suggests that damaging somatic mutations are likely more common than previously appreciated, even in healthy individuals.

As an additional test, we analyzed mosaics in a second cortical region (occipital lobe, 250-300× WGS) and in non-brain tissues (Supplementary Table 9, heart or liver ~40-65× WGS) for two individuals in which these additional tissues were available. We found that 86% of discovered variants, usually those variants with higher VAFs arising earlier during development, were shared in at least two regions, whereas mutations with lower VAFs were often regionally restricted (Fig. 3g-h). Among the 94 sSNVs we identified in the two subjects, one is an exonic missense mutation predicted to be damaging by standard prediction tools (see Methods, Supplementary Table 9), consistent with our estimate that about half of individuals will carry at least one damaging exonic somatic mutation present in a measurable fraction of cells.

### Mutation types evolve in the early embryo and with replication timing

We next assessed the specific base changes in our sSNV dataset. Consistent with other reports of clonal mosaic mutation in humans^13,26,34^, most sSNVs were C>T transitions (48%), with approximately half of those occurring in the context of hypermutable CpG dinucleotides. Interestingly, substitution types evolved across the first four cell divisions with C>T transitions increasing and T>A transitions decreasing (p=7*10^−6^ and p=2*10^−4^, respectively; two-tailed Fisher’s Exact test) in subsequent cell generations (Fig. 4a, Supplementary Fig. 12). CpG C>T mutations, which are believed to be initiated by endogenous spontaneous deamination of 5-methylcytosine, showed the most prominent changes (Fig. 4a, p= 7*10^−6^, Fisher’s Exact test), increasing almost linearly across cell generations.

**Figure 4:**
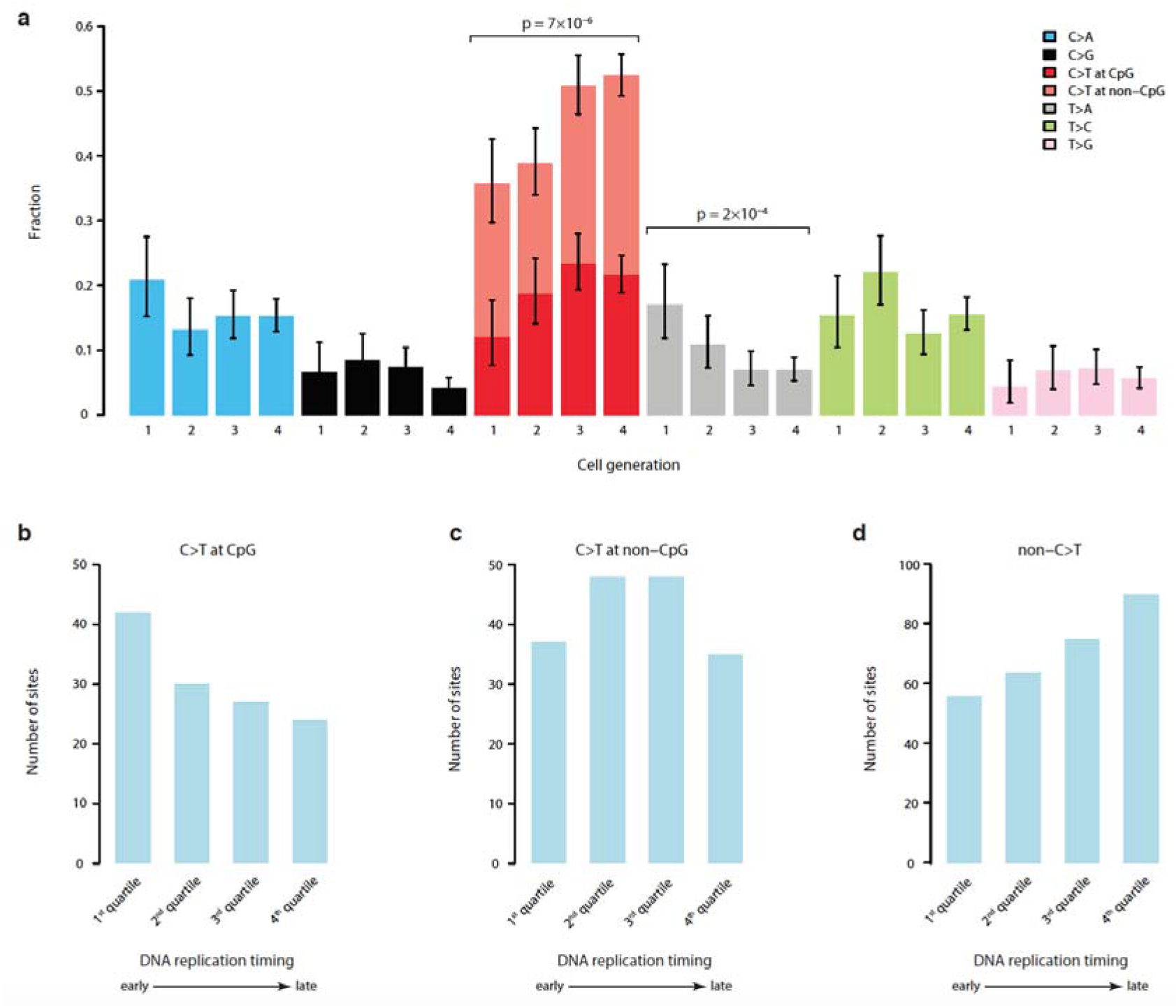
Base substitutions vary with cell generation and replication timing. **a**, There is significant evolution of mutation profiles across the first four cell divisions. Specifically, there is an increase of C>T transitions and decrease of T>A mutations. Error bars indicate 95% CIs calculated with a binomial test. **b,** C>T mutations in CpG dinucleotides tend to occur in earlier replicating genomic regions. **c,** C>T mutations in non-CpG contexts show no replication timing bias. **d,** All other substitution types show a more typical late-replication bias.

Although sSNVs are generally thought to be more prevalent in late-replicating regions of the genome^35,36^, we observed that CpG C>T mutations were more prevalent in the earliest-replicating genomic regions, whereas non-C>T base substitutions increased in later-replicating regions, implying different DNA methylation dynamics during early-embryo development^37^ (Fig. 4b-d).

### Autism brains contain mosaic variants affecting critical brain-active enhancers

Although statistical analysis of WGS studies is challenged by thousands of simultaneous hypotheses that can be tested, the higher rate of mutation in open chromatin that we described above suggested a specific comparison of somatic mutations to a previous study that showed a role of *de novo* germline mutations in neural enhancer sequences in neurodevelopmental disorders^38^. We did not observe enrichment of overall sSNVs and validated mosaic indels in brain-active enhancers in ASD cases compared to controls; however, we did observe significant enrichment when assessing only sequences bearing active enhancer marks in a majority of brain epigenomes available for analysis (from Roadmap Epigenomics^39^), reflecting those regions that are most likely to represent critical enhancers shared across individuals. When restricted to candidates that are recurrent in more than 50% (≥5/8) of available brain epigenomes, the odds ratio of having a mosaic mutation in an enhancer in ASD compared to that in control was 11.9 (95% CI: 1.97-487, p=8*10^−4^, two-tailed Fisher’s Exact test; Fig. 5a, Supplementary Fig. 13-14, Supplementary Table 10). With the Bonferroni correction for testing multiple types of regulatory regions, the p-value is highly significant at p=7*10^−3^. Similar enrichment was not observed in any regulatory elements active in other tissues (Supplementary Fig. 14). While this observation requires confirmation in larger datasets, our data provide a preliminary suggestion that enhancer mutations are not only especially frequent during mitotic cell divisions, but also might contribute to ASD risk in some cases.

Furthermore, genes with transcription start sites within 50kb of these shared brain-active enhancers (≥5 brain epigenomes with active enhancer status in the Roadmap data) with mosaic mutations were enriched for brain-specific expression, implying direct functional relevance of enhancer mutations (p=8*10^−5^, two-tailed Fisher’s Exact test, Fig. 5b, Supplementary Table 11). While non-coding mutations in regulatory regions have been linked to ASD in other studies of peripheral blood DNA^38–43^, our analysis, albeit limited by small sample size, is the first suggestion of such mutations occurring in a mosaic state in brain DNA, suggesting that perturbing the expression of brain-essential genes via brain mosaic mutation in regulatory regions could potentially contribute to ASD risk.

**Figure 5:**
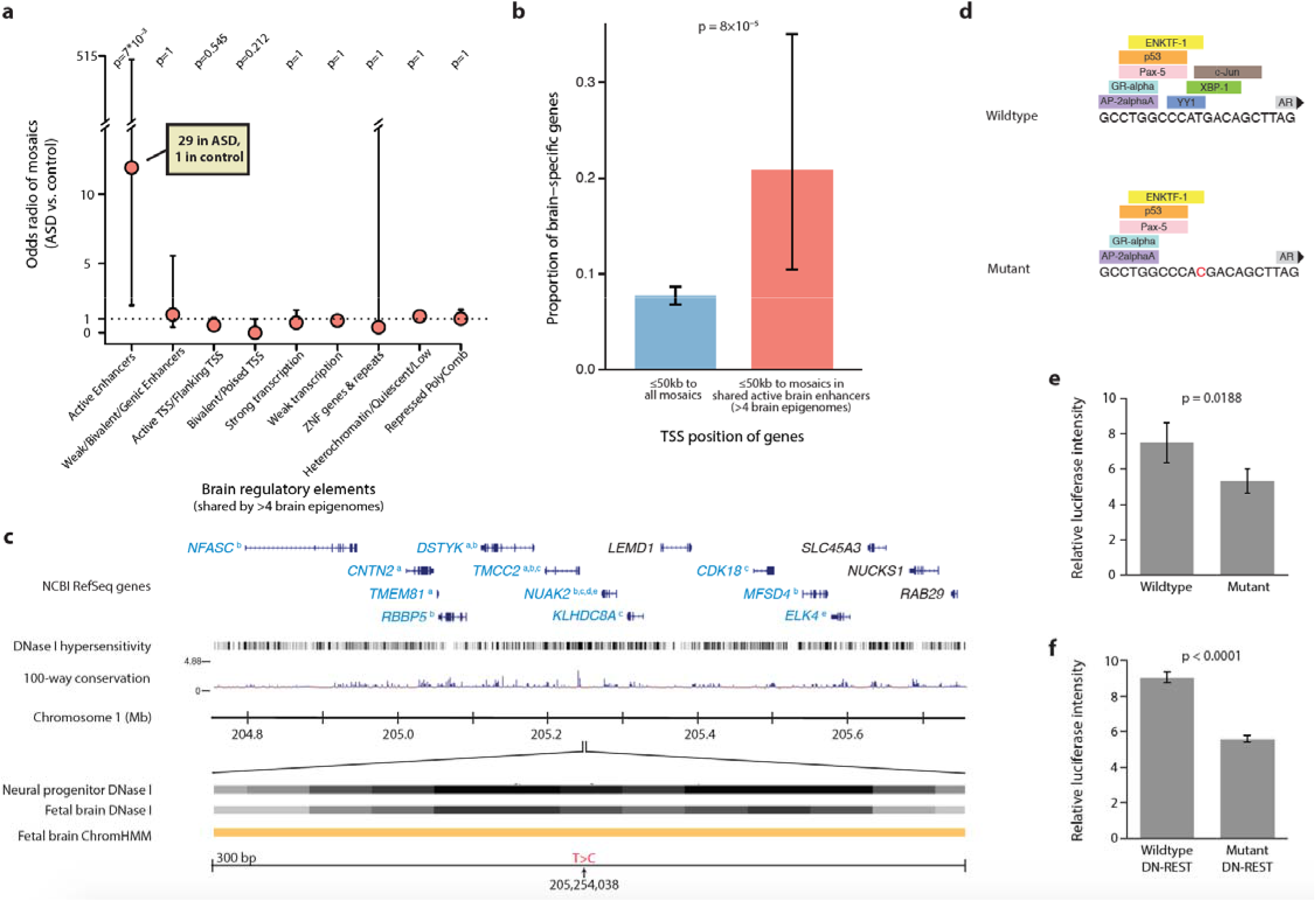
ASD brains contain somatic mutations affecting brain-active enhancers. **a**, Although there is no difference between ASD cases and controls in the burden of mosaic mutations present in all brain-active enhancers from the Roadmap project, ASD cases are enriched for mutations occurring in regions that act as enhancers in the majority of brain epigenomes available for analysis. Odds ratios and error bars (95% CI) were calculated by Fisher’s Exact test, p value was further corrected with Bonferroni correction. Analysis includes sSNVs from PCR-free samples (false positive sites by deep resequencing were excluded) and validated mosaic indels. **b,** Active brain enhancer regions harboring mutations in our dataset are nearby transcription start sites of genes that are enriched for brain-specific expression, compared to genes nearby all mutations in our dataset. Error bars indicate 95% CI calculated with a binomial test. **c,** Example of a sSNV in ASD brain AN06365 located in a brain-active enhancer. Genes in blue font have functional evidence linking their expression to enhancer activity (a = Genotype Tissue Expression^44^, b = Predicted Enhancer Targets^45^, c = Hi-C sequencing data^46^, d = Chromatin Interaction Analysis by Paired-End Tag Sequencing^47^, e = ENCODE data). Orange ChromHMM track represents active enhancer designation. **d,** The mutation is predicted to affect transcription factor binding. **e,** A mutant construct transfected into N2A cells results in reduced enhancer activity by 29% compared to wildtype construct (n=4, two-tailed t-test). **f,** In N2A cells pre-treated with DN-REST to assume a neuronal-like state, the mutant construct reduces enhancer activity by 38% (n=4, two-tailed t-test). Error bars represent 95% CIs.

To investigate the functional significance of putative enhancer mutations, we engineered mutant constructs and assessed their impact on transcriptional activity via a standard luciferase activity assay in cultured neural crest-derived cells for several variants. Case AN06365 has a validated sSNV (present in 5.3% of cells) in an enhancer on chromosome 1 near a number of important genes, some of which have functional data supporting their regulation by the enhancer^44–47^ (Fig. 5c). The single-base substitution identified in this individual is predicted to impact transcription factor binding (Fig. 5d). According to our assay, this mutation has a significant impact on transcriptional activity, resulting in reduced activity in both a neuroblastoma cell line and cells treated with DN-REST in order to produce a more differentiated neuronal state (Fig. 5e-f). Similar results were observed for a second mutation (Supplementary Fig. 15).

## Discussion

In this study we investigated somatic mutations in brain DNA genome-wide using ultra-deep sequencing in a large sample of 74 brains, including 59 brains from patients with ASD. While other studies have performed targeted sequencing of ASD brain DNA^15^, or whole-exome sequencing of brain DNA from other conditions^10^, this represents the largest cohort of ASD brain samples to undergo deep whole-genome analysis. Our study of high-coverage WGS data has revealed a rich landscape of mutational mosaicism within both neurotypical and autistic brains, outlined rates and types of somatic mutation within the brain, and revealed insights into somatic mutation accumulation in the early embryo. We find that the first five cell division produce early somatic mutations in numbers comparable to *de novo* germline mutations, although mosaic mutations will have overall more modest effect sizes given their presence in some but not all brain cells.

Individuals vary substantially in their brain somatic mutation burden, with some brains having only a handful of detectable somatic mutations in a given region and others—even neurotypical individuals—harboring up to several dozen sSNVs present in ≥4-6% of cells, with an elevated mutation rate of sSNVs in exonic and open chromatin regions. Although somatic retrotransposon mobilization has been suggested as a major source of neuronal diversity, sSNVs formed just from the first few cell divisions number ~100 per genome (versus <1 transposon insertion per average genome^48^), in addition to several hundred more sSNVs per genome that arise later in gestation^24^. Given their relative abundance, sSNVs may contribute to inter-individual genetic neural variability more than mobile elements.

Based on our estimated mutation rate per cell division, roughly half of individuals would carry one or more potentially damaging exonic mutations in a substantial fraction of cells (VAF≥1%). At lower VAFs, the number of potentially function-altering somatic variants is expected to be substantially higher. Of note, predicting true deleteriousness is known to be very difficult, such that many mutations predicted as damaging would have functional impact only when homozygous and are likely to be well tolerated especially in a somatic state^49^. Still other mutations may be incompatible with life when homozygous or even when germline heterozygous, meaning that their effects could only be observed in a somatic state. Therefore, although many predicted-damaging mosaic mutations may have subtle effects depending on their distributions, they nonetheless have potential to cause or contribute to a wide number of disease states in many individuals.

Our data also suggest a potential role for mosaic mutations occurring in non-coding regulatory regions in ASD etiology, although the low availability of postmortem brains limits our sensitivity. For example, our sample size was not large enough to identify excess exonic mosaics in autism cases, though this has been documented in much larger exome studies conducted on peripheral DNA from thousands of ASD cases^11–14^. Mosaic mutations in brain-specific enhancer regions are intriguing however, since they represent a mechanism for disrupting gene expression in brain-limited or region-specific ways, in both normal and diseased brains, without disrupting expression in other tissues. Hence, mosaic noncoding mutations represent an attractive candidate mechanism to be involved more broadly in ASD and other neuropsychiatric diseases as well.

## Methods

### Human tissue and DNA samples

Frozen postmortem human brain specimens from 61 ASD cases and 15 neurotypical controls were obtained from the Lieber Institute for Brain Development and the University of Maryland through the NIH Neurobiobank, as well as from Autism BrainNet. All specimens were de-identified and all research was approved by the institutional review board of Boston Children’s Hospital. DNA was extracted from dorsolateral prefrontal cortex where available (or generic cortex in a minority of cases) using lysis buffer from the QIAamp DNA Mini kit (Qiagen) followed by phenol chloroform extraction and isopropanol cleanup. Samples UMB4334, UMB4899, UMB4999, UMB5027, UMB5115, UMB5176, UMB5297, UMB5302, UMB1638, UMB4671, and UMB797 were processed at New York Genome Center using TruSeq Nano DNA library preparation (Illumina) followed by Illumina HiSeq X Ten sequencing to a minimum 200× depth. All remaining samples were processed at Macrogen using TruSeq DNA PCR-Free library preparation (Illumina) followed by minimum 30× sequencing of 7 separate libraries on the Illumina HiSeq X Ten, for a total minimum coverage of 210× per sample. We achieved an average of 251 × depth across all samples, using 150bp paired-end reads. Two samples, UMB5771 and UMB5939, had parental saliva-derived DNA available, and DNA from both parents for these two cases was obtained and sequenced at Macrogen to ~50× depth. Parental DNA was not available for any other samples. Additionally, DNA was extracted from Brodmann Area 17 (occipital lobe) for cases UMB4638 and UMB4643 and sequenced at Macrogen to a minimum 210× depth following PCR-free library preparation. Bulk heart and liver sequencing data, as well as single-cell sequencing data from three individuals (UMB1465, UMB4643, and UMB4638) were previously published by our group and used again in this study^8,17^.

### Mutation calling and filtration

All paired-end FASTQ files were aligned using BWA-MEM version 0.7.8 to the GRCh37/hg19 human reference genome including the hs37d5 decoy sequence from the Broad Institute, following GATK best practices^50,51^. We used MuTect2-PoN^52^ (GATK version: 3.5 nightly-r2016-04-25-g7a7b7cd) to generate a set of PoNs (panel-of-normals) by using 73 individuals other than the sample being analyzed (including both cases and controls), to remove sequencing artifacts and germline variants. Rare variants were further selected by filtering out any variant with a maximum population minor allele frequency >1*10^-15^ in the gnomAD database^53^. Variants within segmental duplication regions or non-diploid regions^16^ were also removed. Low-quality calls tagged “t_lod_fstar,” “str_contraction,” and “triallelic_site” were removed. A minimum VAF of 0.03 was required unless a variant was phasable by Mutect2, which allowed for rescue of variants down to VAF of 0.02; however a threshold of 0.03 was maintained for PCR-based samples. A minimum alternate read depth of 3 reads was required. Only private events among the population were analyzed. An upper VAF threshold of 0.40 was set and heterozygous germline variants were removed. For mosaic indels specifically, variants within RepeatMasker regions (http://www.repeatmasker.org/) and simple repeats regions^54^ were further excluded.

We then used MosaicForecast^16^ to perform read-backed phasing and identify high-confidence mosaics from the candidate call set. Briefly, features likely to be correlated with mosaic detection specificity were selected: mapping quality, base quality, clustering of mutations, read depth, number of mismatches per read, read1/read2 bias, strand bias, base position, read position, trinucleotide context, sequencing cycle, library preparation method, and genotype likelihood. Based on these features a random forest model was trained using phased variants. Further training was conducted using parental WGS data from two cases UMB5771 and UMB5939 as well as single-cell WGS data from three control brains, UMB1465, UMB4643, and UMB4638^8,17^, for which we constructed lineage trees with the sSNVs we identified and assigned variants to different clades, and germline variants were identified as those presenting in multiple conflicting clades^16^. Predicted mosaics were further filtered by removing genomic regions enriched for low-VAF variants and by removing variants with unusually high sequencing depth that also occurred in regions marked as copy number variants (CNVs) by Meerkat^55^. Following all training and filtration, we identified 2166 putative mosaic sSNVs (Supplementary Table 2). Two ASD sample, MSSM007 and UMB5308, were eliminated from the study at this stage due to very high noise suggestive of sample contamination, leaving 59 ASD cases with high-quality sequencing data.

### Prediction of pathogenicity scores

Pathogenicity prediction scores were calculated for functional mosaic and germline variants using a modified version of a previously described pipeline^18^. The pipeline uses 12 different prediction tools (SIFT, LRT, MutationTaster, MutationAssesor, FATHMM, Provean, MetaSVM, MetaLR, M-CAP, MutPred, Eigen, and CADD) and classifies variants as follows: Nsyn = missense variant with ≥5 benign predictions or 0 damaging predictions; NsynD1 = damaging missense variant with ≥1 damaging prediction and <5 benign predictions; NsynD2 = damaging missense variant with ≥4 damaging predictions and <5 benign predictions; NsynD4 = damaging missense ≥5 damaging predictions and <5 benign predictions and GERP >2 (CADD >15 || DANN >0.9 || EIGAN >0.9 || REVEL >0.9); LOF-1 = stopgain/frameshift; LOF-2 = canonical splicing (intronic +/− 1 and 2 bases); LOF-3 = exonic splicing sites +/− 2bp or intronic splicing region (+/-3-15bp) plus splicing impact prediction; LOF-4 = Other sites with large splicing prediction; LOF-5 = Stop-loss, likely benign mutations in splicing regions, extended splicing, not predicted to cause change, and GERP <2. For germline mutation analysis, mutations were identified in genes present in the Simons Foundation Autism Research Initiative (SFARI) database of ASD-relevant genes (http://gene.sfari.org/) with a score of 1 (high confidence involvement in ASD) or 2 (strong candidate for involvement in ASD). Gene constraint was calculated with pLI scores^56^ and with missense and synonymous Z scores^57^. Loss-of-function was also assessed with LOFTEE analysis^58^. Genes were also screened through the Online Inheritance in Man (OMIM) database of genes with relevance to any human disease (http://www.omim.org/).

### Amplicon resequencing validation

Targeted validation was attempted on 208 mosaic sSNVs and all called indels. Additional validation was conducted on called exonic sSNVs that were ultimately excluded from the dataset due to low VAF in PCR-based samples or presence in gnomAD database (Supplementary Table 6). Validation candidates were selected based on potential functional significance, ability to design PCR primers, and representative diversity of VAFs and genomic loci. Multiple sets of PCR primers were designed for each variant and synthesized with Ion Torrent adapters P and A, with barcodes added for unique identification. PCR amplification was performed using Phusion HotStart II DNA Polymerase (Thermo) as described by the manufacturer, with 20-25 cycles of amplification. Reactions were pooled and purified with AMPure XP technology (Agencourt), then sequenced on the Ion Torrent Personal Genome Machine using the Ion 530 chip with 400bp reads, reaching an average coverage of 92,000 reads per variant, amongst sSNV reactions that yielded mappable reads.

Following demultiplexing and trimming, reads were mapped using BWA-MEM and locally realigned using GATK. High-quality reference and mutant reads were then counted using mpileup and variants with successful PCR reactions resulting in usable reads were then classified as validated true mosaics or homozygous reference with variant not present. Any ambiguous variants, including variants in which there was discordance between sequencing from different PCR primers, were conservatively assigned a designation of homozygous reference. Validation success rates were calculated as the number of true mosaics divided by the sum of true mosaics and homozygous reference, excluding variants from brains UMB1465, UMB4643, and UMB4638 as validation in these brains was conducted on an alternative DNA source as none of the originally sequenced DNA remained. Weighted averaging across PCR and PCR-free variant validation was used to determine a comprehensive validation rate of 90%. Five variants from UMB5771 and UMB5939 were also re-sequenced in parent DNA, which confirmed a mosaic state in the offspring and homozygous reference in parents.

### Epigenetic covariates of mosaic mutations

Candidate mosaic mutations were annotated with ANNOVAR^59^ (Version: 2017-07-17) to calculate the observed density of putative mosaics in different regions. Rare SNPs (MAF<0.01) from 15,708 whole genomes of unrelated individuals in the Genome Aggregation Database (gnomAD)^57^ were annotated with ANNOVAR and used to calculate the expected density of mutations in different regions. DNAse I hypersensitive regions for different tissues were downloaded from http://egg2.wustl.edu/roadmap/data/byFileType/peaks/consolidated/broadPeak/DNase/, and the DNAseI-accessible Regulatory Regions (FDR0.01) were used to calculate the in- and out-of-region density of putative mosaics. We merged DHS regions in different tissues profiled by the Roadmap Epigenomics Project to obtain a single set of DHS regions. Chromatin states in different tissues and cell lines predicted by Hidden Markov Model v1.10 using 18 states (6 marks, 98 epigenomes) were downloaded from https://egg2.wustl.edu/roadmap/data/byFileType/chromhmmSegmentations/ChmmModels/core_K27ac/jointModel/final/^60^. Mutations in ASD patients versus control samples in different regulatory regions were compared using a two-tailed Fisher’s exact test. A Bonferroni correction for multiple hypothesis testing was implemented for nine comparisons as specified below. Roadmap epigenomes were separated for analysis based on their tissue of origin. States were classified as follows: 9_EnhA1 and 10_EnhA2 = active enhancers, 7_EnhG1, 8_EnhG2, 11_EnhWk and 15_EnhBiv = weak/bivalent/genic enhancers, 1_TssA, 2_TssFlnk, 3_TssFlunkU and 4_TssFlnkD = active TSS/flanking TSS, 14_TssBiv = bivalent/poised TSS, 5_Tx = strong transcription, 6_TxWk = weak transcription, 12_ ZNF/Rpts = ZNF genes and repeats, 13_Het and 18_Quies = heterochromatin/quiescent/low, 16_TssBiv and 17_ReprPCWk = repressed polycomb.

### Simulation of mosaic mutations and calculation of sensitivity

300× WGS data for NA12878 (Genome in Bottle, downloaded from ftp://ftp-trace.ncbi.nlm.nih.gov/giab) was downsampled to 250× using SAMtools^61^. High-confidence SNP calls for individual NA12878 were downloaded from ftp://ftp-trace.ncbi.nih.gov/giab/ftp/release/NA12878_HG001/NISTv2.18/^62^. Simulated mosaic mutations with different VAFs were generated in the 250× BAM file by converting bases supporting the alternate alleles of high-confidence heterozygous SNPs to reference bases at several binomial sampling probabilities. Simulated sites with expected VAF of 0.01, 0.02, 0.03, 0.04, 0.05, 0.08, 0.1, 0.15, 0.2, 0.25, 0.3, 0.32, 0.34, 0.36, 0.38, 0.4, 0.45 and 0.5 were generated and used to calculate sensitivities. 95% C.I.s for sensitivity at each VAF were calculated using a binomial test.

### Symmetric vs. asymmetric cell contribution analysis

Ion Torrent amplicon resequencing for 96 germline heterozygous mutations revealed that VAFs were over-dispersed compared to a binomial distribution (Supplementary Fig. 3), likely due to noise induced by PCR amplification as part of resequencing. We fit the VAF distribution with a beta-binomial model to capture the over-dispersion 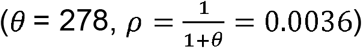. An R package VGAM^63^ (v1.1-1) was used to fit the beta-binomial model. It has previously been reported that the two ancestor cells of the blood lineage give rise to offspring asymmetrically at approximately a 2:1 ratio^25^. Here we used Ion Torrent-validated mosaics from diploid chromosomes with a similar model to measure potential asymmetrical cell contributions to the brain during early embryonic development. Briefly, we let *α*_1_ and 1 – *α*_1_ be the fraction of brain cells deriving from each of the two cells created by the first division of the brain ancestor cell. A contribution parameter value of *α*_1_ = 0.5 means the first two cells contributed equally to the brain, while a non-0.5 value means the cell contribution is asymmetrical. Given a specific *α*_1_ it is possible to calculate the expected VAF for mutations acquired at different branches of the early phylogeny. Assuming the mutation rate per cell generation is constant (i.e., the two cell divisions from the 2^nd^ cell generation have the same mutation rate), we compute the likelihood of a mosaic arising on a specific branch by multiplying the estimated sensitivity for detecting mosaics at the expected branch VAF with the over-dispersion beta-binomial likelihood of the mosaic VAF measured by the deep Ion Torrent sequencing. The log likelihoods for all sites were then summed over all branches to estimate the log likelihood of a specific *α*_1_. We fit *α*_1_ by maximizing the log likelihood over *α* ∈ [0.5,1] using a grid search with step size = 0.0002. A likelihood ratio test was used to compare the asymmetrical model to the symmetrical model = 0.5), which favored the model with unequal cell contribution during the 1^st^ cell generation (p =3*10^−4^). A 95% C.I. for *α*_1_ (0.555, 0.597, Supplementary Fig. 10) was constructed using the likelihood ratio (all values of *α*_1_ for which the likelihood drops off by no more than 1.92 units).

To examine the potential AF dispersion problem in our 250× WGS data, we randomly extracted 25,000 phasable germline sites with 0.4-0.6 AF (by MuTect2) and plotted the AF distribution profile. No AF over-dispersion was found compared with binomial sampling (Supplementary Fig. 3).

### Mutation rate estimation and assignment of mutations to cell generations

To estimate per-generation mutation rates, we used an expectation-maximization algorithm similar to that described by Ju *et al*.^25^. Briefly, the mutation rate *v_g_* for all cell generations was considered to be identical at the beginning, and the probability of a mutation *j* belonging to cell generation *g* (expectation step) is

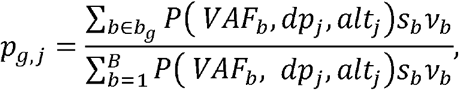

where *P(VAF_b_, dpj,αltj)* is the binomial probability of observing *alt_j_* successes (alternate allele supporting reads) in *dpj* trials (total reads) with probability of success *VAF_b_. B* denotes the total number of branches for the 1^st^-5^th^ cell generations; *v_b_* is the mutation rate for branch *b; b_g_* is the set of all branches belonging to generation g; and *s_b_* denotes the sensitivity for detecting mosaics on branch *b*. We assumed the same mutation rate for all branches in a specific generation, and symmetrical contributions of cells to the embryo. The mutation rate for cell generations g was then updated as the sum of *p_g,j_* across all mosaics (maximization step):

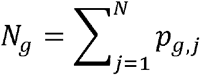

The two steps were iterated until convergence.

To obtain upper and lower bounds for mutation rate per cell generation, we ran several different bootstrap simulations over mosaics from the 63 PCR-free samples. Bootstrap resamplings (sampling 63 brains each time) were performed 1,000 times to estimate the distribution of observed mutations per cell division (Supplementary Fig. 7), and for each cell division the total number of mosaics was obtained by dividing the observed number of mosaics by the VAF-specific detection sensitivities. A 95% C.I. was computed by calculating 2.5-97.5% percentiles.

The mutation rate for the 4^th^ cell generation and the mutation rate for the 5^th^ cell generation were added up to give an estimate of the mutation rate for 4+ cell generations, and the mutation rates per cell division for the 1^st^, 2^nd^, 3^rd^ and 4+ cell generations were estimated to be 3.37, 2.51,2.28, and 2.85 respectively. Using our WGS data, the probability of each mutation belonging to each cell generation was obtained, and all 1641 putative mutations from PCR-free samples and Ion Torrent-validated PCR-based samples were then assigned to different cell generations using maximum likelihood values. 182 mutations were assigned to the 1^st^ cell generation, 250 to the 2^nd^ cell generation, 392 to the 3^rd^ cell generation, and 817 to the 4^th^ cell generation and beyond (Supplementary Table 8). We also compared mutation-to-generation assignments determined by deep WGS bulk data to those calculated from single-cell data for three previously analyzed individuals^8,17^ and found reasonable agreement between the two methods (Fig. 3b and Supplementary Fig. 11). Our cell generation assignments did not change appreciably when based on asymmetric vs. symmetric models of cell division (98.6% in concordance, Supplementary Table 8).

### Estimation of total mutations and exonic mutations per individual

There are 31 cell divisions (2^0^ + 2^1^ + 2^2^ + 2^3^ + 2^4^) in the first five cell generations of early embryonic development. Mutations per cell division for the 1^st^-5^th^ cell generations were bootstrapped from the values we generated in the section above, and the total number of mutations was calculated by adding up all mutations from the 1^st^-5^th^ cell generations. The process was repeated 10,000 times to estimate the total number of mutations per individual. We calculated number of exonic mutations using our data that 2.2% of called sSNVs were exonic. Reported values were obtained by simulating binomial sampling on the total number of mutations for each individual. A 95% C.I. was computed by calculating 2.5-97.5% percentiles.

### Comparison of mosaics in brain and non-brain tissues

Mosaic mutations were called from WGS data for two different brain regions (PFC and BA17, representing occipital lobe) in two individuals (UMB4643 and UMB4638) using the same pipeline described above. All putative mosaic mutations from each region were visually inspected by SAMtools mpileup across different tissues, including the two brain regions with ~250× depth of coverage and one non-brain tissue (liver or heart) with ~50× depth of coverage. A mutation was considered to be absent from the tissue if there were no alternative allele-supporting reads observed in that tissue.

### Comparison of mutational signatures between earlier and later mutations

We downloaded all 96-dimensional mutational signatures from PCAWG^64^. To avoid over-fitting, we extracted the two most common clock-like signatures (Signature S1 and Signature S5) as well as a signature highly related with sequencing artifacts (Signature S18) from the PCAWG signatures, and deconstructed mutational signatures for the mosaic mutations using the R package deconstructSigs^65^. We observed a trend toward an increase in signature S1 across the 1^st^-4^th^ cell generations (Supplementary Fig. 12), which is believed to be caused by an endogenous mutational process initiated by spontaneous deamination of 5-methylcytosine^66^ Mutation profiles from different cell generations were compared using a two-tailed Fisher’s exact test.

### Comparison of DNA replication timing between mutations from different cell generations

We extracted locus-specific DNA replication timing for all putative mosaic mutations using AsymTools (http://software.broadinstitute.org/cancer/cga/AsymTools)^67^. The sSNVs were then classified into four categories according to DNA replication time quartiles.

### Evaluation of gene expression level with TSSs nearby shared brain active enhancers

Transcription start sites (TSSs) of coding genes were extracted from Gencode^68^ v19 annotations on GRCh37. Genes with their TSSs overlapping regions within 50kb upstream or downstream of sSNVs from PCR-free samples were extracted. Tissue-specific expression derived for a total of 53 tissues and cell types were downloaded from GTEx project^69^. We used the expression table from GTEx v.7 (gene median transcripts per million [TPM] per tissue) and brain-specific genes were defined as: 1. Genes with median expression level across brains three times higher than the median expression level across all tissues; 2. The tissue with highest median expression level is a brain tissue. A two-tailed Fisher’s Exact Test was applied to compare types of genes (brain-specific genes or others) nearby sSNVs (genes with TSSs adjacent to mosaics in shared brain active enhancers versus genes with TSSs adjacent to all mosaic sites).

### Luciferase assays for assessment of enhancer activity

We selected 17 sSNVs identified in brain-active enhancers and attempted cloning and site-directed mutagenesis (New England Biolabs) to recreate the mutations. Mutagenesis was successful for 11 mutations. Wildtype and mutant constructs were then cloned into luciferase vector pGL4.25 (Promega). Luciferase plasmids were transfected into N2A cells, along with an internal control plasmid (phRL-TK(Int-), Promega) and dominant-negative REST (DN-REST)^70^ or GFP expression plasmids using Polyfect (Qiagen). Luciferase activities were measured 24 hours later. Three wildtype enhancer constructs had technically successful assays with significant difference from negative control, and among these two showed significant difference between wildtype and mutant. All experiments were performed with n=4 and averaged across replicates.

## Acknowledgements

Human tissue was obtained from the NIH NeuroBioBank at the University of Maryland, the Lieber Institute for Brain Development, Oxford University Brain Bank, and Autism BrainNet. We thank the donors and their families for their invaluable contribution to the advancement of science. We also thank R. Sean Hill, Jennifer Partlow, Wayne Bainter, and the Research Computing group at Harvard Medical School for assistance. RER is supported by the Stuart H.Q. and Victoria Quan Fellowship in Neurobiology, and by the Harvard/MIT MD-PhD program (T32GM007753). Y.D., M.K., M.A.S., D.C.G., and P.J.P. are supported by a grant from the NIMH (U01MH106883, P50MH106933) and the Harvard Ludwig Center. C.A.W. is supported by the Manton Center for Orphan Disease Research, the Allen Discovery Center program through The Paul G. Allen Frontiers Group, grants from the NINDS (R01 NS032457 and U01 MH106883), and grant U01MH106883 from the NIMH. C.A.W. is an Investigator of the Howard Hughes Medical Institute. Data were generated as part of the Brain Somatic Mosaicism Network (BSMN) Consortium, supported by: U01MH106874, U01MH106876, U01MH106882, U01MH106883, U01MH106883, U01MH106884, U01MH106891, U01MH106891, U01MH106891, U01MH106892, U01MH106893, U01MH108898 awarded to: Nenad Sestan (Yale University), Flora Vaccarino (Yale University), Fred Gage (Salk Institute for Biological Studies), Christopher Walsh (Boston Children’s Hospital), Peter J. Park (Harvard University), Jonathan Pevsner (Kennedy Krieger Institute), Andrew Chess (Icahn School of Medicine at Mount Sinai), John V. Moran (University of Michigan), Daniel Weinberger (Lieber Institute for Brain Development), and Joseph Gleeson (University of California, San Diego). The content of this paper is solely the responsibility of the authors and does not necessarily represent the official views of the National Institute of General Medical Sciences or the National Institutes of Health.

## Members of The Brain Somatic Mosaicism Network

**Boston Children’s Hospital:** Christopher Walsh, Javier Ganz, Mollie Woodworth, Pengpeng Li, Rachel Rodin, Robert Hill, Sara Bizzotto, Zinan Zhou

**Harvard University:** Alice Lee, Alison Barton, Alissa D’Gama, Alon Galor, Anna Chung, Craig Bohrson, Daniel Kwon, Doga Gulhan, Elaine Lim, Euncheon Lim, Giorgio Melloni, Isidro Cortes, Jake Lee, Jia Wang, Joe Luquette, Lixing Yang, Maxwell Sherman, Michael Coulter, Michael Lodato, Peter Park, Rebeca Monroy, Semin Lee, Sonia Kim, Soo Lee, Yanmei Dou

**Icahn School of Medicine at Mt. Sinai:** Andrew Chess, Attila Gulyas-Kovacs, Chaggai Rosenbluh, Schahram Akbarian

**Kennedy Krieger Institute:** Ben Langmead, Jeremy Thorpe, Jonathan Pevsner, Sean Cho

**Lieber Institute for Brain Development:** Andrew Jaffe, Apua Paquola, Daniel Weinberger, Jennifer Erwin, Jooheon Shin, Richard Straub, Rujuta Narurkar

**Mayo Clinic:** Alexej Abyzov, Taejeong Bae

**NIMH:** Anjene Addington, David Panchision, Doug Meinecke, Geetha Senthil, Lora Bingaman, Tara Dutka, Thomas Lehner

**Rockefeller University:** Laura Saucedo-Cuevas, Tara Conniff

**Sage Bionetworks:** Kenneth Daily, Mette Peters

**Salk Institute for Biological Studies:** Fred Gage, Meiyan Wang, Patrick Reed, Sara Linker

**Stanford University:** Alex Urban, Bo Zhou, Xiaowei Zhu

**Universitat Pompeu Fabra:** Aitor Serres, David Juan, Inna Povolotskaya, Irene Lobon, Manuel Solis, Raquel Garcia, Tomas Marques-Bonet

**University of California, Los Angeles:** Gary Mathern

**University of California, San Diego:** Eric Courchesne, Jing Gu, Joseph Gleeson, Laurel Ball, Renee George, Tiziano Pramparo

**University of Michigan:** Diane A. Flasch, Trenton J. Frisbie, Jeffrey M. Kidd, John B. Moldovan, John V. Moran, Kenneth Y. Kwan, Ryan E. Mills, Sarah Emery, Weichen Zhou, Yifan Wang

**University of Virginia:** Aakrosh Ratan, Mike McConnell

**Yale University:** Flora Vaccarino, Gianfilippo Coppola, Jessica Lennington, Liana Fasching, Nenad Sestan, Sirisha Pochareddy

## Author contributions

R.E.R., Y.D., P.J.P., and C.A.W. conceptualized study. R.E.R., A.M.D., and S.N.K. generated whole-genome sequencing data. Y.D. led bioinformatic analysis with assistance from M.K. for variant identification and from M.A.S., L.J.L., C.L.B., and D.C.G for technical issues. R.E.R., L.R. and R.N.D. performed targeted variant validation. L.R. and K.M.G. performed cloning and luciferase assay experiments. R.E.R., Y.D., P.J.P., and C.A.W. wrote the manuscript.

## Code availability statement

Custom code is available from the authors by request.

## Data availability statement

Whole-genome sequencing data will be available from the National Institute of Mental Health Data Archive (DOI: 10.15154/1503337).

## Supplementary Figures

**Supplementary Figure 1:** Analysis pipeline.

**Supplementary Figure 2:** Sensitivity of mosaic detection measured with simulated mosaics with different VAFs.

**Supplementary Figure 3:** Over-dispersion of VAF measured by Ion Torrent deep resequencing.

**Supplementary Figure 4:** Distribution of mosaic indel lengths.

**Supplementary Figure 5:** Number of exonic mosaics per individual.

**Supplementary Figure 6:** An Ion Torrent-validated deleterious missense mosaic mutation in the ASD risk gene *CACNA1A.*

**Supplementary Figure 7:** Bootstrap simulations of mutation rate per cell division.

**Supplementary Figure 8:** Consistently high validation rate of candidate mosaics from PCR-free samples with different VAFs, measured by Ion Torrent deep re-sequencing.

**Supplementary Figure 9:** Evaluation of putative mosaics with high allele fraction (VAF>0.2) in single cell sequencing data.

**Supplementary Figure 10:** Asymmetric cell contribution likelihood test.

**Supplementary Figure 11:** Assignment of mosaic mutations to different cell generations based on asymmetric cell model.

**Supplementary Figure 12**: Evolving spectra of somatic mutations in generations 1-4+ of embryonic cell divisions and trend in signatures across cell divisions.

**Supplementary Figure 13:** Brain-active enhancer enrichment remains present when PCR-based samples are included in analysis.

**Supplementary Figure 14:** Mosaics in active enhancer regions were not enriched in non-brain tissues.

**Supplementary Figure 15:** Brain-active enhancer mutation in case UMB4671.

## Supplementary Tables

**Supplementary Table 1:** ASD cases and controls used in this study. Brain region indicates left hemisphere unless otherwise noted. PMI = Post-Mortem Interval; SUDEP = Sudden Unexplained Death in Epilepsy; ADI-R = Autism Diagnostic Interview – Revised, with category A representing social interaction (cutoff 10), category B representing verbal or nonverbal communication (cutoff 8 for verbal, 7 for nonverbal), category C representing restricted, repetitive, and stereotyped behaviors (cutoff 3), and category D representing developmental abnormalities present before age 36 months (cutoff 1).

**Supplementary Table 2:** All 2166 mosaic mutations identified in this study.

**Supplementary Table 3:** Validation results for PCR-free and PCR-based variants.

**Supplementary Table 4:** List of 96 germline heterozygous sites covered in deep resequencing.

**Supplementary Table 5:** Validation results for indels.

**Supplementary Table 6:** All exonic mutations.

**Supplementary Table 7:** Rare damaging germline heterozygous variants in autism risk genes earning score of 1 or 2 in SFARI gene list. All listed variants have maximum population minor allele frequency <0.001 in Kaviar, 1000 Genomes, EVS6500, ExAC-nonpsych, and gnomAD.

**Supplementary Table 8:** Maximum likelihood estimation of cell generations in both symmetrical and asymmetrical models for 1641 sSNVs from PCR-free samples and validated PCR-based mutations.

**Supplementary Table 9:** Distribution of sSNVs detected in WGS from PFC, occipital lobe, and non-brain tissue of two individuals UMB4643 and UMB4638.

**Supplementary Table 10:** List of sSNVs and somatic indels occurring in brain-active enhancers, summary data for Roadmap epigenomes, and additional enhancer mutations excluded from dataset.

**Supplementary Table 11:** Genotype-tissue expression data for genes adjacent to mosaic mutations.

## References

1 Lynch M. Rate, molecular spectrum, and consequences of human mutation. Proc Natl Acad Sci U SA 107, 961–968, doi:10.1073/pnas.0912629107 (2010).

2 D’Gama, A. M. et al. Mammalian target of rapamycin pathway mutations cause hemimegalencephaly and focal cortical dysplasia. Ann Neurol 77, 720–725, doi: 10.1002/ana.24357 (2015).

3 Lim, J. S. et al. Somatic Mutations in TSC1 and TSC2 Cause Focal Cortical Dysplasia. Am J Hum Genet 100, 454–472, doi:10.1016/j.ajhg.2017.01.030 (2017).

4 Nakashima, M. et al. Somatic Mutations in the MTOR gene cause focal cortical dysplasia type IIb. Ann Neurol 78, 375–386, doi:10.1002/ana.24444 (2015).

5 Erickson, R. P. Recent advances in the study of somatic mosaicism and diseases other than cancer. Curr Opin Genet Dev 26, 73–78, doi:10.1016/j.gde.2014.06.001 (2014).

6 Insel, T. R. Brain somatic mutations: the dark matter of psychiatric genetics? Mol Psychiatry 19, 156–158, doi:10.1038/mp.2013.168 (2014).

7 McConnell, M. J. et al. Intersection of diverse neuronal genomes and neuropsychiatric disease: The Brain Somatic Mosaicism Network. Science 356, doi:10.1126/science.aal1641 (2017).

8 Lodato, M. A. et al. Aging and neurodegeneration are associated with increased mutations in single human neurons. Science 359, 555–559, doi:10.1126/science.aao4426 (2018).

9 Keogh, M. J. et al. High prevalence of focal and multi-focal somatic genetic variants in the human brain. Nat Commun 9, 4257, doi:10.1038/s41467-018-06331-w (2018).

10 Wei, W. et al. Frequency and signature of somatic variants in 1461 human brain exomes. Genet Med 21, 904–912, doi:10.1038/s41436-018-0274-3 (2019).

11 Dou, Y. et al. Postzygotic single-nucleotide mosaicisms contribute to the etiology of autism spectrum disorder and autistic traits and the origin of mutations. Hum Mutat 38, 1002–1013, doi:10.1002/humu.23255 (2017).

12 Freed, D. & Pevsner, J. The Contribution of Mosaic Variants to Autism Spectrum Disorder. PLoS Genet 12, e1006245, doi:10.1371/journal.pgen.1006245 (2016).

13 Lim, E. T. et al. Rates, distribution and implications of postzygotic mosaic mutations in autism spectrum disorder. Nat Neurosci 20, 1217–1224, doi:10.1038/nn.4598 (2017).

14 Krupp, D. R. et al. Exonic Mosaic Mutations Contribute Risk for Autism Spectrum Disorder. Am J Hum Genet 101, 369–390, doi:10.1016/j.ajhg.2017.07.016 (2017).

15 D’Gama, A. M. et al. Targeted DNA Sequencing from Autism Spectrum Disorder Brains Implicates Multiple Genetic Mechanisms. Neuron 88, 910–917, doi:10.1016/j.neuron.2015.11.009 (2015).

16 Dou, Y. et al. Accurate detection of mosaic variants in sequencing data without matched controls. Nat Biotechnol, doi:10.1038/s41587-019-0368-8 (2020).

17 Lodato, M. A. et al. Somatic mutation in single human neurons tracks developmental and transcriptional history. Science 350, 94–98, doi:10.1126/science.aab1785 (2015).

18 Doan, R. N. et al. Recessive gene disruptions in autism spectrum disorder. Nat Genet 51, 1092–1098, doi:10.1038/s41588-019-0433-8 (2019).

19 Damaj, L. et al. CACNA1A haploinsufficiency causes cognitive impairment, autism and epileptic encephalopathy with mild cerebellar symptoms. Eur J Hum Genet 23, 1505–1512, doi:10.1038/ejhg.2015.21 (2015).

20 Epi, K. C. De Novo Mutations in SLC1A2 and CACNA1A Are Important Causes of Epileptic Encephalopathies. Am J Hum Genet 99, 287–298, doi:10.1016/j.ajhg.2016.06.003 (2016).

21 Mercer, T. R. et al. DNase I-hypersensitive exons colocalize with promoters and distal regulatory elements. Nat Genet 45, 852–859, doi:10.1038/ng.2677 (2013).

22 Thurman, R. E. et al. The accessible chromatin landscape of the human genome. Nature 489, 75–82, doi:10.1038/nature11232 (2012).

23 Polak, P. et al. Reduced local mutation density in regulatory DNA of cancer genomes is linked to DNA repair. Nat Biotechnol 32, 71–75, doi:10.1038/nbt.2778 (2014).

24 Ye, A. Y. et al. A model for postzygotic mosaicisms quantifies the allele fraction drift, mutation rate, and contribution to de novo mutations. Genome Res 28, 943–951, doi:10.1101/gr.230003.117 (2018).

25 Ju, Y. S. et al. Somatic mutations reveal asymmetric cellular dynamics in the early human embryo. Nature 543, 714–718, doi:10.1038/nature21703 (2017).

26 Bae, T. et al. Different mutational rates and mechanisms in human cells at pregastrulation and neurogenesis. Science 359, 550–555, doi:10.1126/science.aan8690 (2018).

27 Rahbari, R. et al. Timing, rates and spectra of human germline mutation. Nat Genet 48, 126–133, doi:10.1038/ng.3469 (2016).

28 Wong, C. C. et al. Non-invasive imaging of human embryos before embryonic genome activation predicts development to the blastocyst stage. Nat Biotechnol 28, 1115–1121, doi:10.1038/nbt.1686 (2010).

29 Kiessling, A. A. et al. Genome-wide microarray evidence that 8-cell human blastomeres overexpress cell cycle drivers and under-express checkpoints. J Assist Reprod Genet 27, 265–276, doi: 10.1007/s10815-010-9407-6 (2010).

30 Gonzalez-Marin, C., Gosalvez, J. & Roy, R. Types, causes, detection and repair of DNA fragmentation in animal and human sperm cells. Int J Mol Sci 13, 14026–14052, doi:10.3390/ijms131114026 (2012).

31 Russell, L. B. & Russell, W. L. Spontaneous mutations recovered as mosaics in the mouse specific-locus test. Proc Natl Acad Sci U S A 93, 13072–13077, doi:10.1073/pnas.93.23.13072 (1996).

32 Neale, B. M. et al. Patterns and rates of exonic de novo mutations in autism spectrum disorders. Nature 435, 242–245, doi:10.1038/nature11011 (2012).

33 Kryukov, G. V., Pennacchio, L. A. & Sunyaev, S. R. Most rare missense alleles are deleterious in humans: implications for complex disease and association studies. Am J Hum Genet 80, 727–739, doi: 10.1086/513473 (2007).

34 Blokzijl, F. et al. Tissue-specific mutation accumulation in human adult stem cells during life. Nature 538, 260–264, doi:10.1038/nature19768 (2016).

35 Chen, C. L. et al. Impact of replication timing on non-CpG and CpG substitution rates in mammalian genomes. Genome Res 20, 447–457, doi:10.1101/gr.098947.109 (2010).

36 Koren, A. et al. Differential relationship of DNA replication timing to different forms of human mutation and variation. Am J Hum Genets 91, 1033–1040, doi:10.1016/j.ajhg.2012.10.018 (2012).

37 Seisenberger, S. et al. The dynamics of genome-wide DNA methylation reprogramming in mouse primordial germ cells. Mol Cell 48, 849–862, doi:10.1016/j.molcel.2012.11.001 (2012).

38 Short, P. J. et al. De novo mutations in regulatory elements in neurodevelopmental disorders. Nature 555, 611–616, doi:10.1038/nature25983 (2018).

39 Williams, S. M. et al. An integrative analysis of non-coding regulatory DNA variations associated with autism spectrum disorder. Mol Psychiatry 24, 1707–1719, doi:10.1038/s41380-018-0049-x (2019).

40 Zhou, J. et al. Whole-genome deep-learning analysis identifies contribution of noncoding mutations to autism risk. Nat Genet 51, 973–980, doi:10.1038/s41588-019-0420-0 (2019).

41 An, J. Y. et al. Genome-wide de novo risk score implicates promoter variation in autism spectrum disorder. Science 362, doi:10.1126/science.aat6576 (2018).

42 Turner, T. N. et al. Genomic Patterns of De Novo Mutation in Simplex Autism. Cell 171, 710–722 e712, doi:10.1016/j.cell.2017.08.047 (2017).

43 Turner, T. N. et al. Genome Sequencing of Autism-Affected Families Reveals Disruption of Putative Noncoding Regulatory DNA. Am J Hum Genet 98, 58–74, doi:10.1016/j.ajhg.2015.11.023 (2016).

44 Consortium, G. T. Human genomics. The Genotype-Tissue Expression (GTEx) pilot analysis: multitissue gene regulation in humans. Science 348, 648–660, doi:10.1126/science.1262110 (2015).

45 He, B, Chen, C., Teng, L. & Tan, K. Global view of enhancer-promoter interactome in human cells. Proc Natl Acad Sci U S A 111, E2191–2199, doi:10.1073/pnas.1320308111 (2014).

46 Jin, F. et al. A high-resolution map of the three-dimensional chromatin interactome in human cells. Nature 503, 290–294, doi:10.1038/nature12644 (2013).

47 Li, G. et al. Extensive promoter-centered chromatin interactions provide a topological basis for transcription regulation. Cell 148, 84–98, doi:10.1016/j.cell.2011.12.014 (2012).

48 Evrony, G. D. et al. Single-neuron sequencing analysis of L1 retrotransposition and somatic mutation in the human brain. Cell 151, 483–496, doi:10.1016/j.cell.2012.09.035 (2012).

49 Miosge, L. A. et al. Comparison of predicted and actual consequences of missense mutations. Proc Natl Acad Sci USA 112, E5189–5198, doi:10.1073/pnas.1511585112 (2015).

50 Genovese, G., Handsaker, R. E., Li, H., Kenny, E. E. & McCarroll, S. A. Mapping the human reference genome’s missing sequence by three-way admixture in Latino genomes. Am J Hum Genet 93, 411–421, doi:10.1016/j.ajhg.2013.07.002 (2013).

51 McKenna, A. et al. The Genome Analysis Toolkit: a MapReduce framework for analyzing nextgeneration DNA sequencing data. Genome Research 20, 1297–1303 (2010).

52 Cibulskis, K. et al. Sensitive detection of somatic point mutations in impure and heterogeneous cancer samples. Nat Biotechnol 31, 213–219, doi:10.1038/nbt.2514 (2013).

53 Karczewski, K. J. et al. Variation across 141,456 human exomes and genomes reveals the spectrum of loss-of-funotion intolerance across human protein-coding genes. bioRxiv, 531210, doi:10.1101/531210 (2019).

54 Benson, G. Tandem repeats finder: a program to analyze DNA sequences. Nucleic Acids Res 27, 573–580, doi: 10.1093/nar/27.2.573 (1999).

55 Yang, L. et al. Diverse mechanisms of somatic structural variations in human cancer genomes. Cell 153, 919–929, doi:10.1016/j.cell.2013.04.010 (2013).

56 Samocha, K. E. et al. A framework for the interpretation of de novo mutation in human disease. Nat Genet 46, 944–950, doi:10.1038/ng.3050 (2014).

57 Lek, M. et al. Analysis of protein-coding genetic variation in 60,706 humans. Nature 536, 285–291, doi:10.1038/nature19057 (2016).

58 MacArthur, D. G. et al. A systematic survey of loss-of-function variants in human protein-coding genes. Science 335, 823–828, doi:10.1126/science.1215040 (2012).

59 Wang, K, Li, M. & Hakonarson, H. ANNOVAR: functional annotation of genetic variants from high-throughput sequencing data. Nucleic Acids Res 38, e164, doi:10.1093/nar/gkq603 (2010).

60 Roadmap Epigenomics, C. et al. Integrative analysis of 111 reference human epigenomes. Nature 518, 317–330, doi:10.1038/nature14248 (2015).

61 Li, H. et al. The Sequence Alignment/Map format and SAMtools. Bioinformatics 25, 2078–2079, doi:10.1093/bioinformatics/btp352 (2009).

62 Rimmer, A. et al. Integrating mapping-, assembly- and haplotype-based approaches for calling variants in clinical sequencing applications. Nat Genet 46, 912–918, doi:10.1038/ng.3036 (2014).

63 Yee, T. W. Vector Generalized Linear and Additive Models: With an Implementation in R. (2015).

64 Alexandrov, L. et al. The Repertoire of Mutational Signatures in Human Cancer. bioRxiv, doi:10.1101/322859 (2018).

65 Rosenthal, R., McGranahan, N., Herrero, J., Taylor, B. S. & Swanton, C. DeconstructSigs: delineating mutational processes in single tumors distinguishes DNA repair deficiencies and patterns of carcinoma evolution. Genome Biol 17, 31, doi:10.1186/s13059-016-0893-4 (2016).

66 Alexandrov, L. B. et al. Signatures of mutational processes in human cancer. Nature 500, 415–421, doi:10.1038/nature12477 (2013).

67 Haradhvala, N. J. et al. Mutational Strand Asymmetries in Cancer Genomes Reveal Mechanisms of DNA Damage and Repair. Cell 164, 538–549, doi:10.1016/j.cell.2015.12.050 (2016).

68 Frankish, A. et al. GENCODE reference annotation for the human and mouse genomes. Nucleic Acids Res 47, D766–D773, doi:10.1093/nar/gky955 (2019).

69 Consortium, G. T. et al. Genetic effects on gene expression across human tissues. Nature 550, 204–213, doi:10.1038/nature24277 (2017).

70 Chong, J. A. et al. REST: a mammalian silencer protein that restricts sodium channel gene expression to neurons. Cell 80, 949–957, doi:10.1016/0092-8674(95)90298-8 (1995).

