## Supplementary Figures for "The Landscape of Mutational Mosaicism in Autistic and Normal Human Cerebral Cortex"

**Supplementary Figure 1**

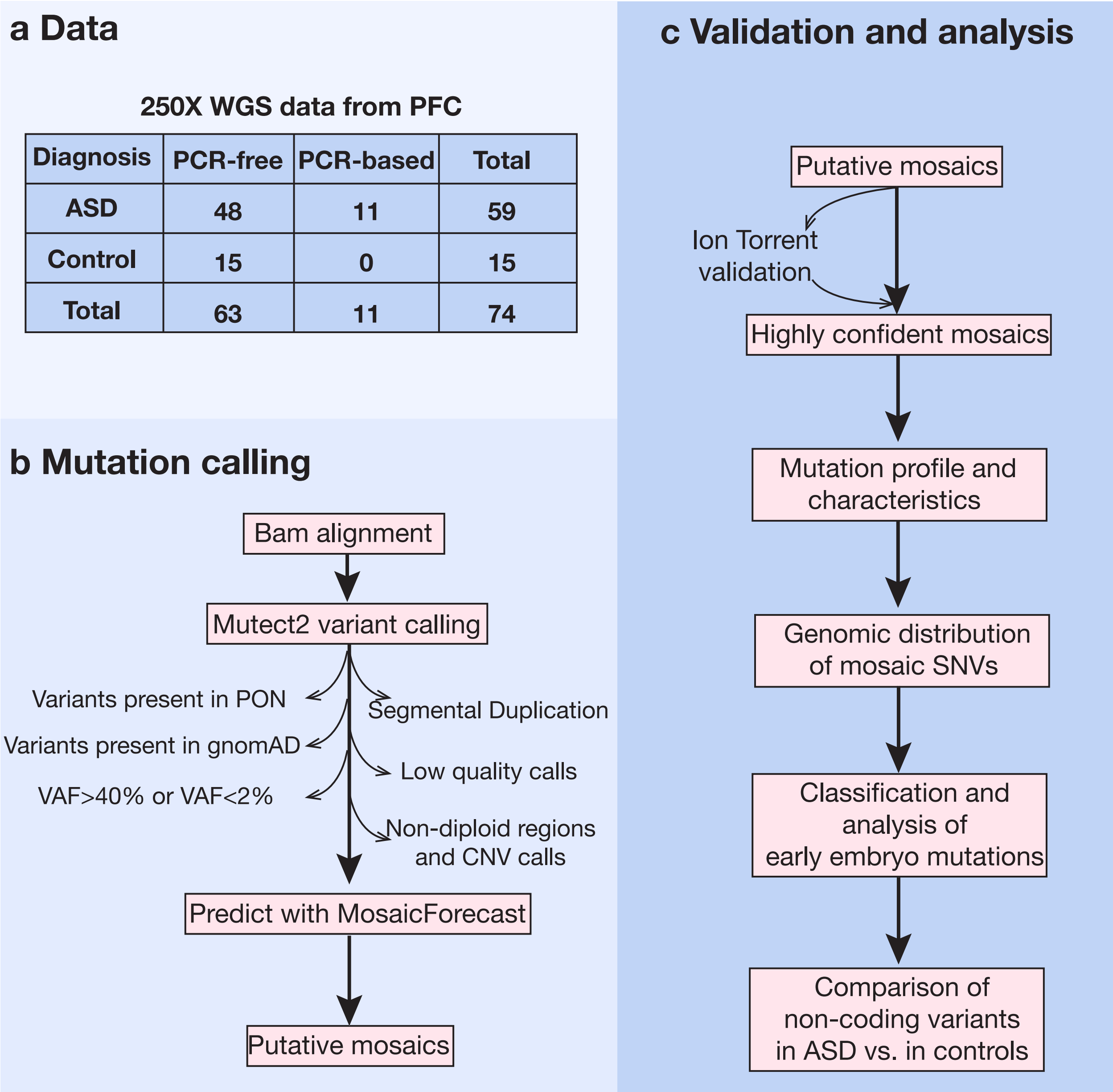


**Analysis pipeline**. **a**, Overview of the largest ultra-deep whole-genome sequencing data set of autism brain-derived DNA, including 59 ASD cases and 15 neurotypical controls. **b**, Following alignment of sequencing reads to the human reference genome, germline and putative somatic variants were called using MuTect2 against a panel of normals. To identify high-confidence mosaic mutations, mutations were filtered based on VAF, population minor allele fraction, quality, and genomic context, and assessed with our novel machine learning tool MosaicForecast. Clustered mutations and mutations overlapping copy number variants were eliminated, yielding a highly confident set of mosaic mutations. **c**, A representative set of mutations was validated using targeted sequencing, and downstream analyses of mutation characteristics, spectra, genomic regions, and burdens were performed. Our validation showed that mosaics from PCR-free samples had a very high validation rate and all were used in all following analyses; however, mosaics from PCR-based samples had a lower validation rate and were therefore excluded from most downstream analyses except where specified.

**Supplementary Figure 2**

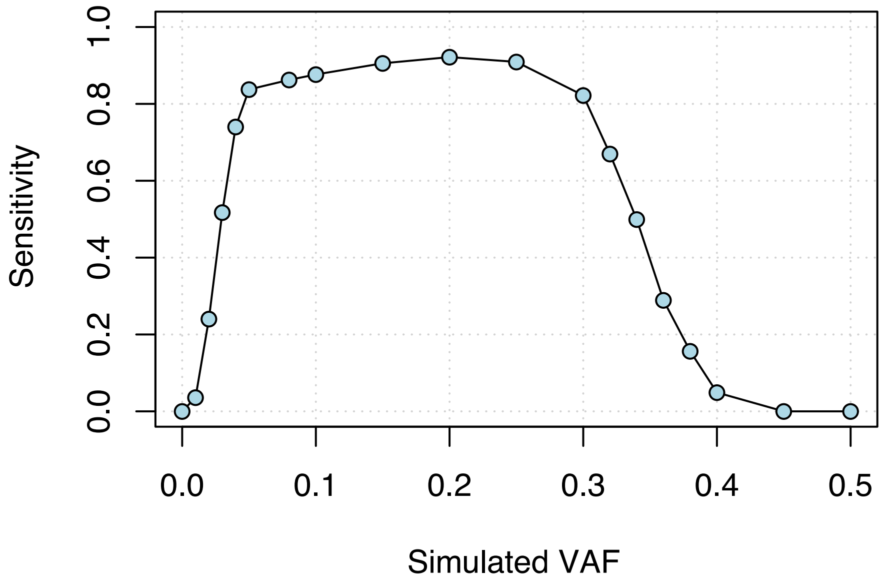


**Sensitivity of mosaic detection measured with simulated mosaics with different VAFs in non-repeat regions.** Based on simulated data, our mutation calling algorithm is highly sensitive for variants with VAF between ~5% and ~33%.

#### Supplementary Figure 3

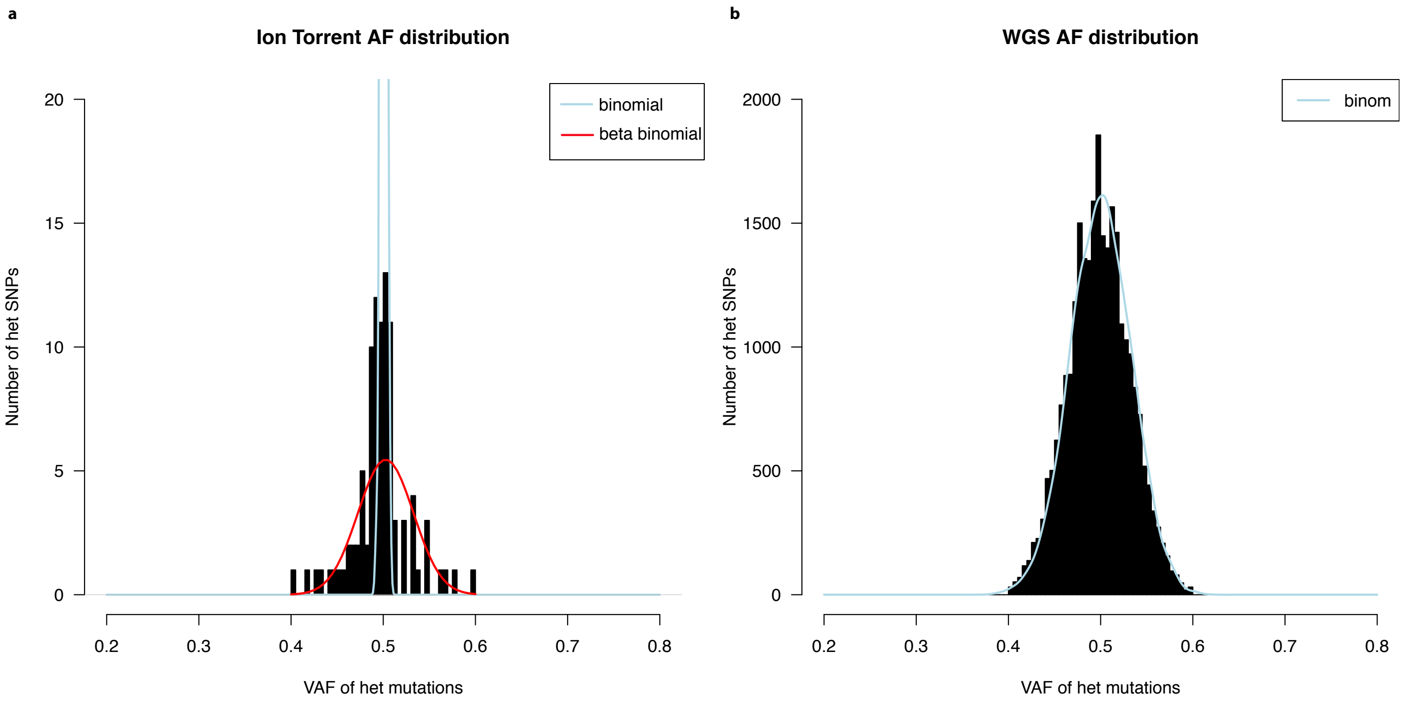


**Over-dispersion of VAF measured by Ion Torrent deep re-sequencing. a**, VAFs from Ion Torrent amplicon resequencing for germline heterozygous mutations were over-dispersed compared with the binomial distribution. Asymmetrical cell contribution parameters were corrected for overdispersion as described in methods, and the red curve is the density curve of the beta-binomial distribution after correction. **b**, Germline heterozygous variants called from WGS data were not over-dispersed compared with binomial distribution.

**Supplementary Figure 4**

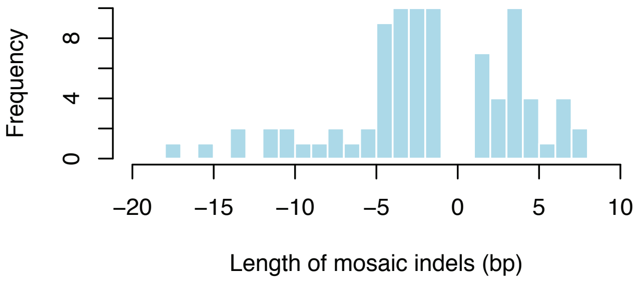


**Length distribution of mosaic indels identified and validated in this study.** Although many 1bp indels were called, they were technically difficult to distinguish from artifact in validation experiments.

**Supplementary Figure 5**

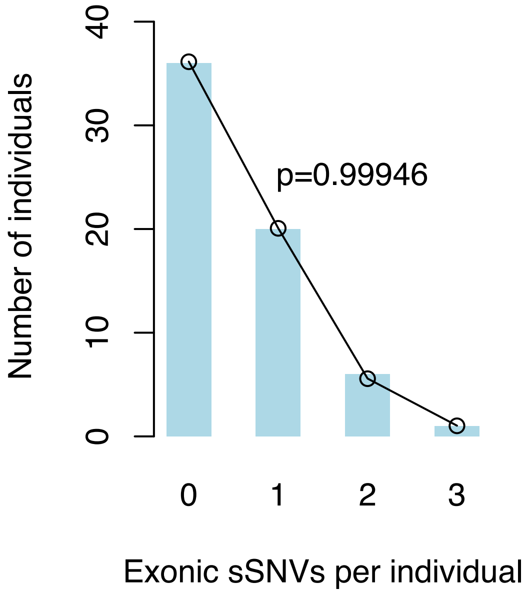


**Distribution of exonic sSNVs per subject.** The occurrence of sSNVs per individual conforms to a Poisson distribution (chi-square test).

**Supplementary Figure 6**

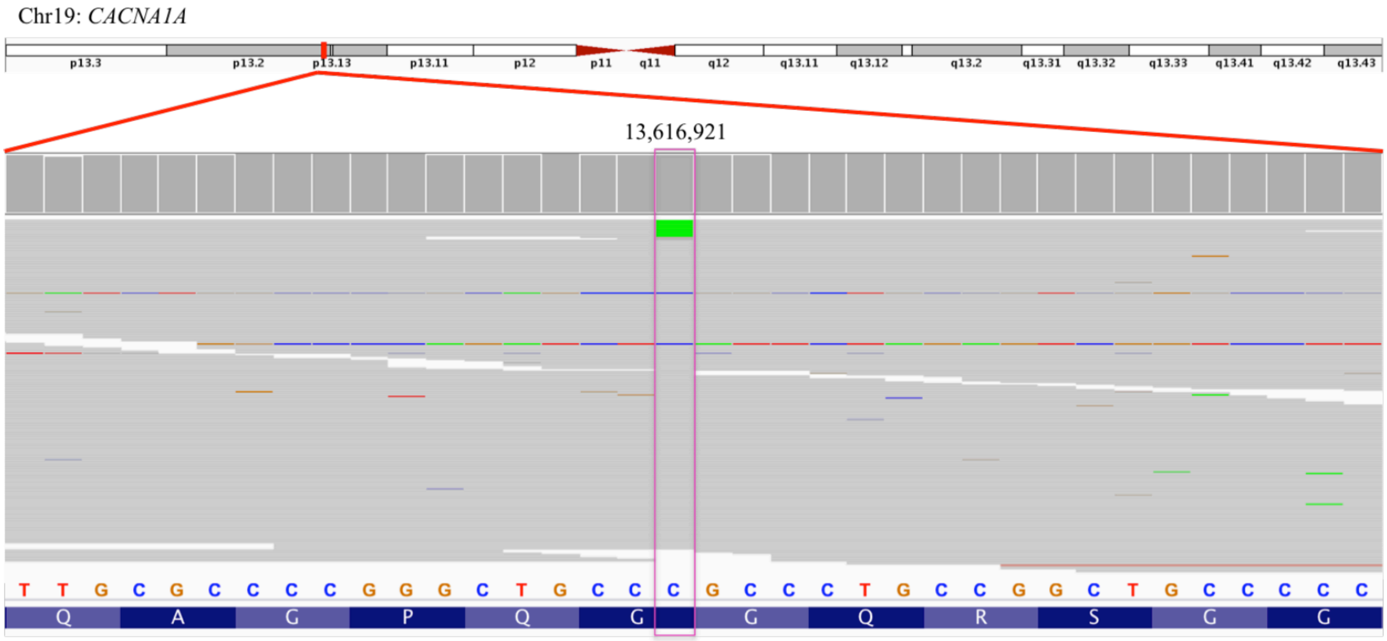


**An Ion Torrent-validated deleterious missense mosaic mutation in the ASD risk gene *CACNA1A*.** A deleterious missense C>A change in the ASD risk gene *CACNA1A* was called in 5.2% of sequencing reads in case UMB1174. Targeted validation of this region in a total of 93,000 reads confirmed a VAF of 5.0%, meaning that this mutation is present in ~10% of cells.

#### Supplementary Figure 7

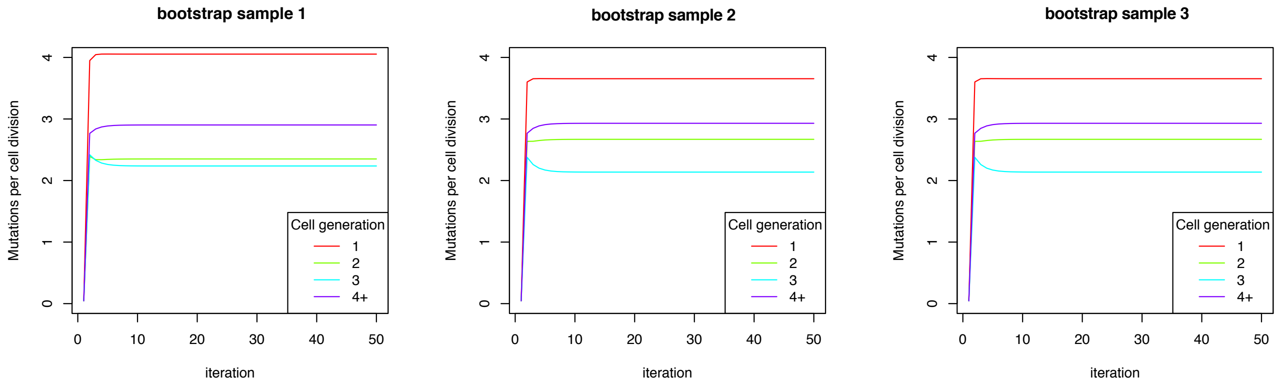


**Bootstrap simulations of mutation rate per cell division consistently show elevated mutation rate in the first cell cycle.** Each panel represents one bootstrap resampling (sampling 63 brains each time). Mutation rates were calculated using an EM algorithm as described in the Methods (50 iterations for convergence).

**Supplementary Figure 8**

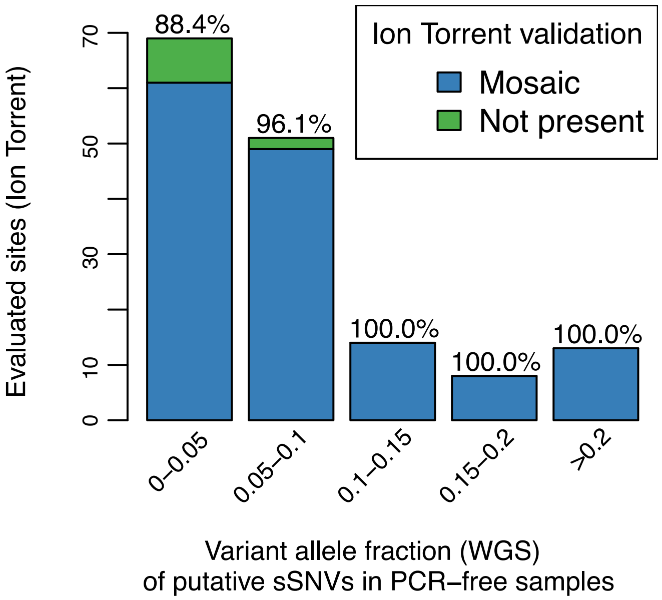


**Consistently high validation rate of candidate mosaics from PCR-free samples with different VAFs, measured by Ion Torrent deep re-sequencing.** sSNVs with >20% VAFs have a 100% validation rate, indicating that the high-AF mosaics are unlikely to be germline variants.

### Supplementary Figure 9

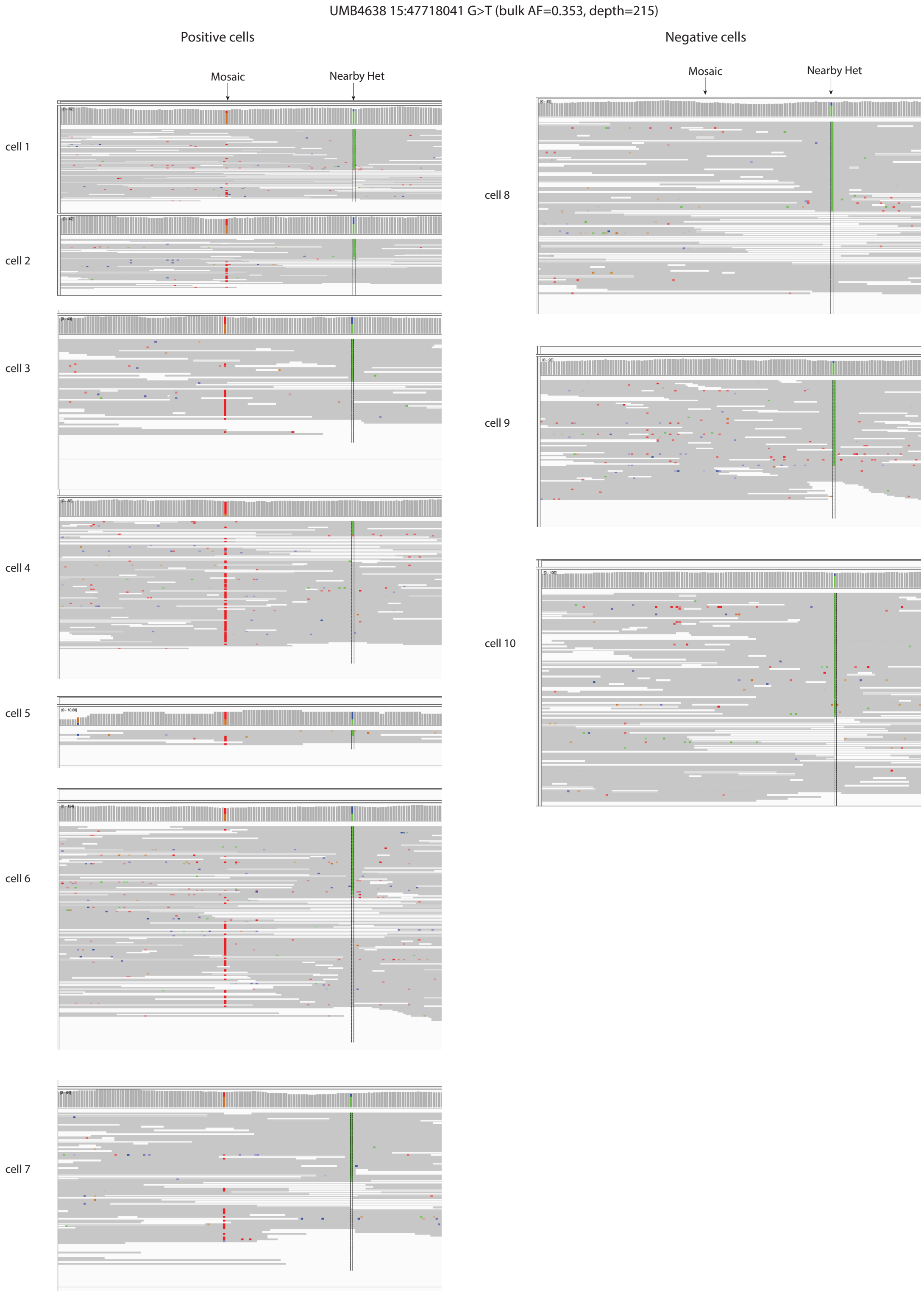

**(continued)**

UMB4638 3: 55954310 G>T (bulk AF=0.215, read depth=260)

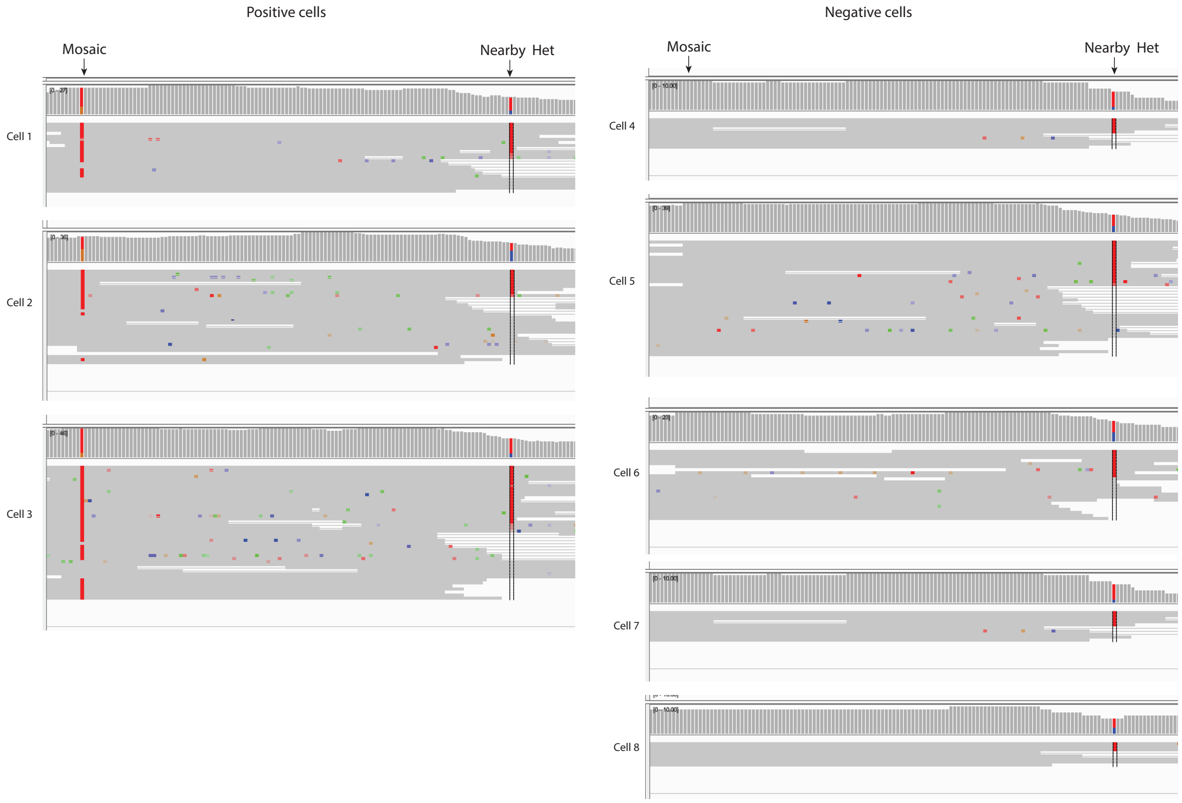

**(continued)**

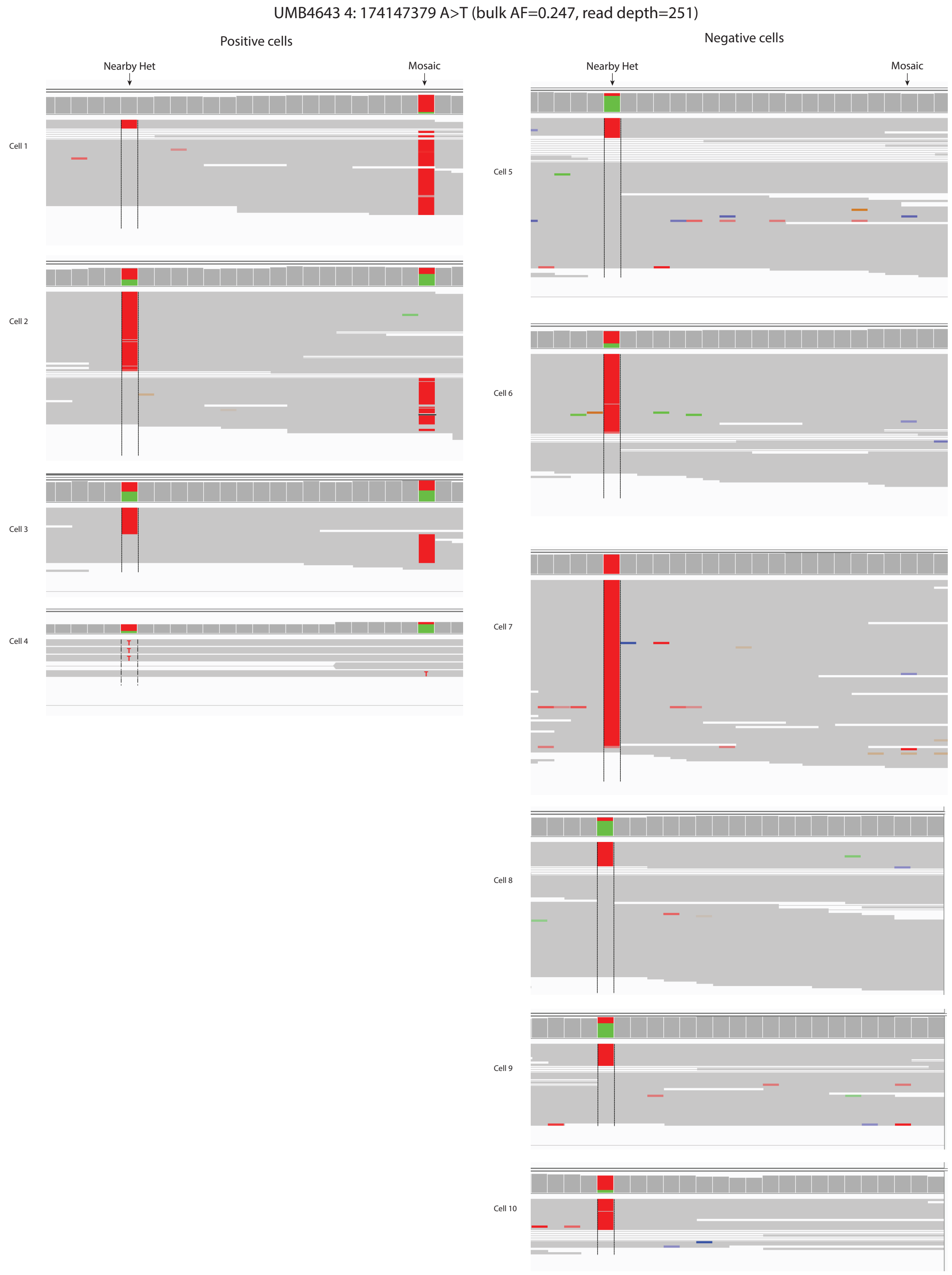
UMB4643 4: 174147379 A>T (bulk AF=0.247, read depth=251)
Positive cells
Negative cells
Nearby Het
Mosaic
Nearby Het
Mosaic
Cell 1
Cell 2
Cell 3
Cell 4
Cell 5
Cell 6
Cell 7
Cell 8
Cell 9
Cell 10

**(continued)**

UMB4643 11:95795713 A>G (bulk AF=0.219, read depth=247)

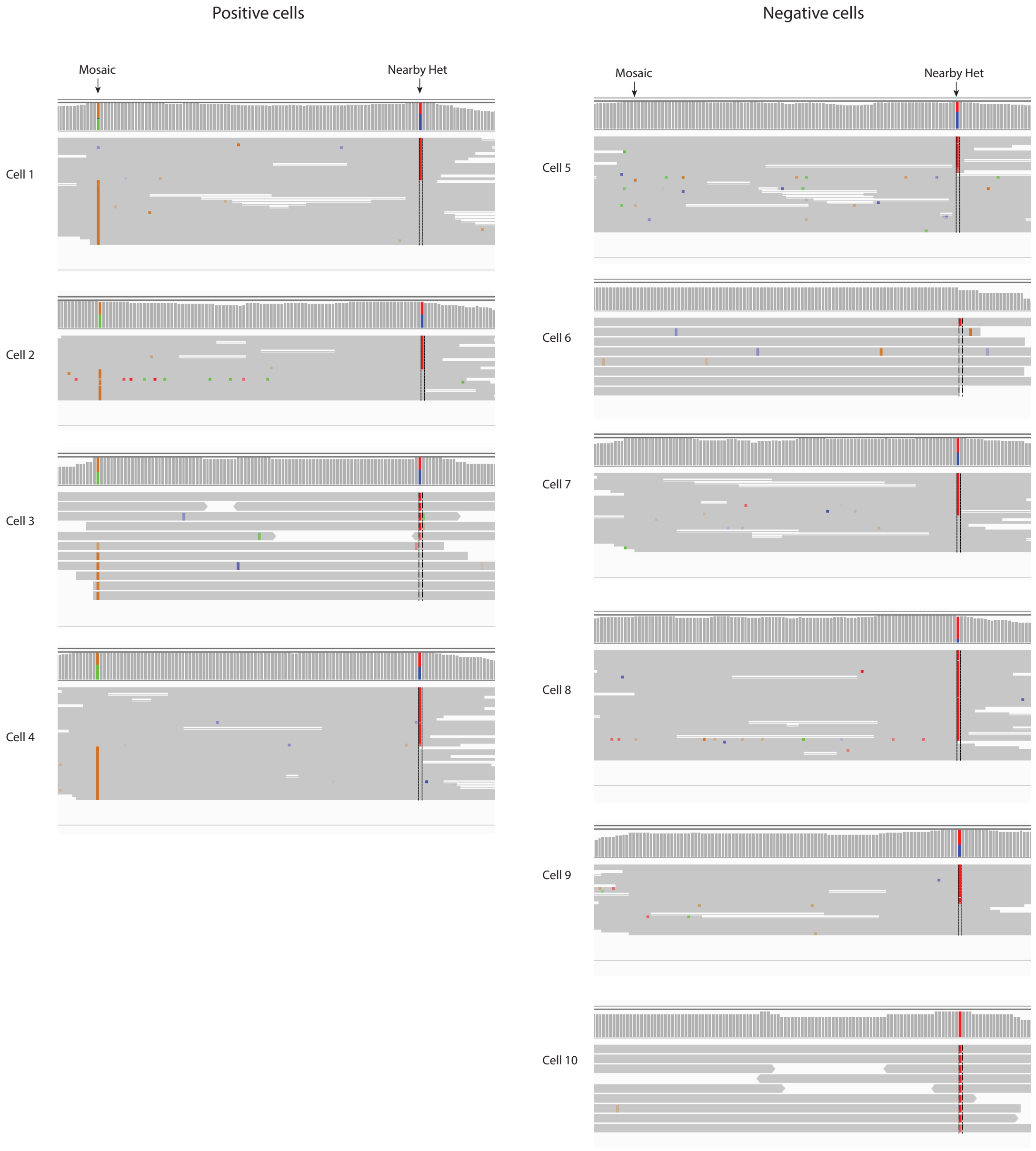

**(continued)**

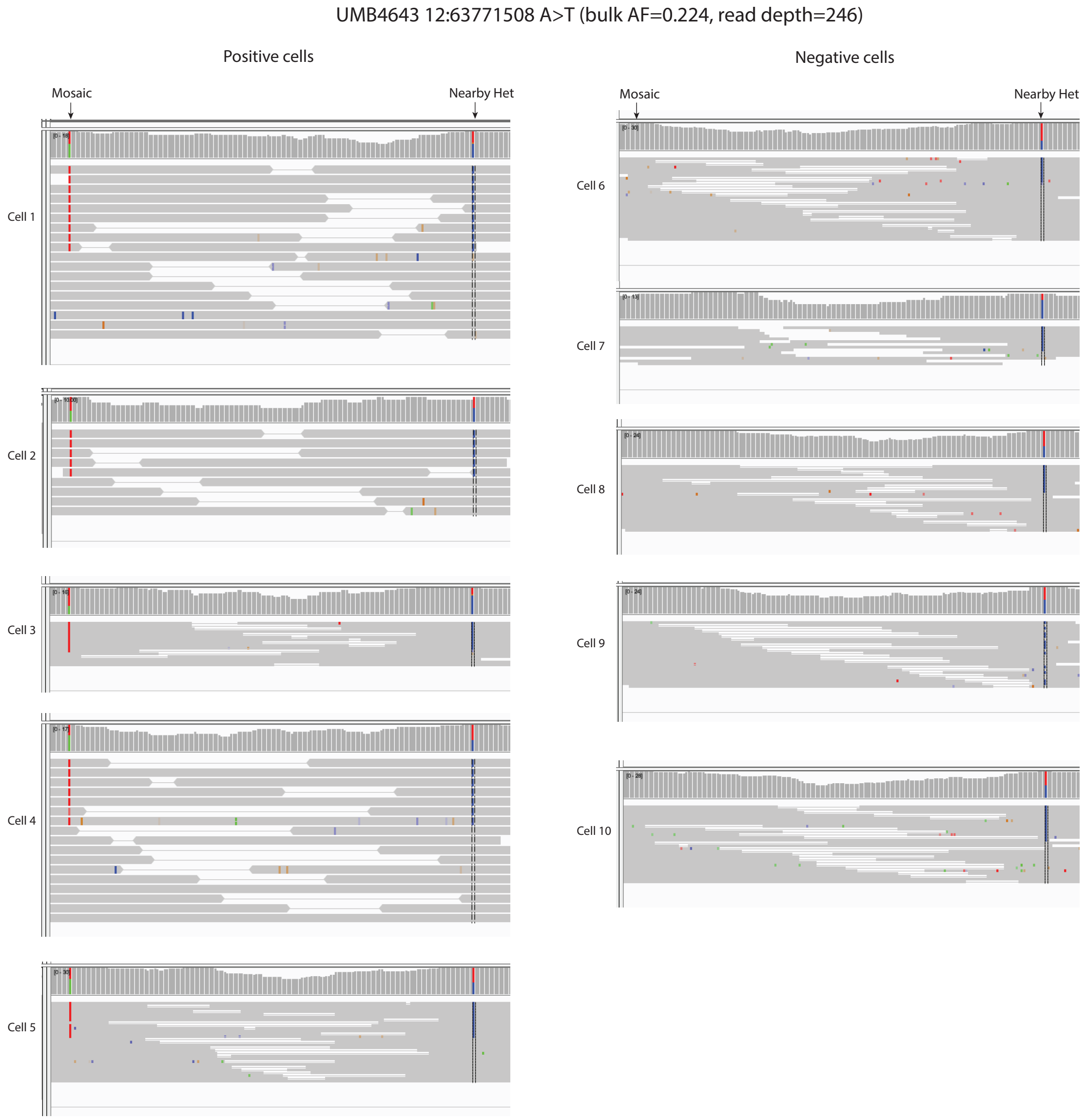


**Evaluation of putative mosaics with high allele fraction (VAF>0.2) in single cell sequencing data.** All six variants with VAF >0.2 and a nearby heterozygous variant from two individuals 4643 and 4638 were evaluated in single cell data. The variant 11:69807919 A>T from individual 4643 was not shown here because it has low coverage across all single cells. The two individuals 4643 and 4638 both have 10 single cells with WGS sequencing data available. High allele fraction mosaics with VAF>0.2 called by MosaicForecast are present in about half of the single cells as shown in this figure. For the phasable sites with at least one nearby germline variant, the two alleles of the nearby germline variant still exist in most non-mutant cells, indicating that the absence of mutation in these cells is not due to allelic dropout. In the cells carrying the mosaic mutation, the mutation is completely linked with one haplotype of the heterozygous mutation (hap=2), indicating that these are heterozygous cells. Thus, about half of cells are reference homozygous and about half of cells are heterozygous according to the single cell data, demonstrating that they are true mosaic variants.

**Supplementary Figure 10**

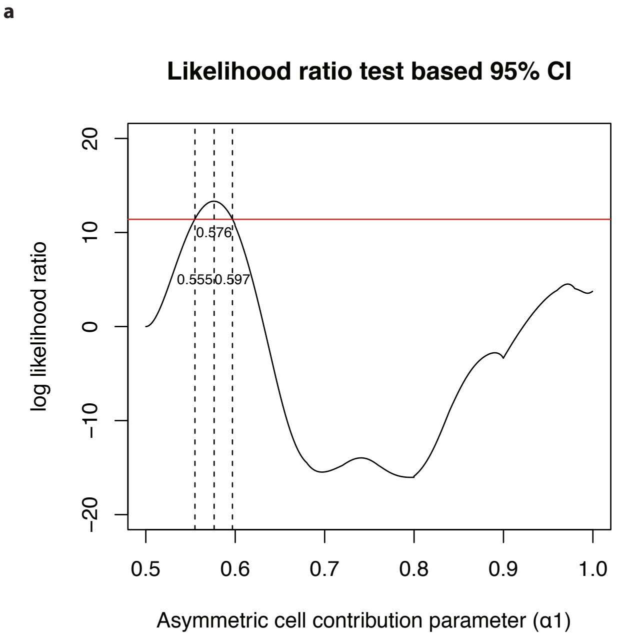


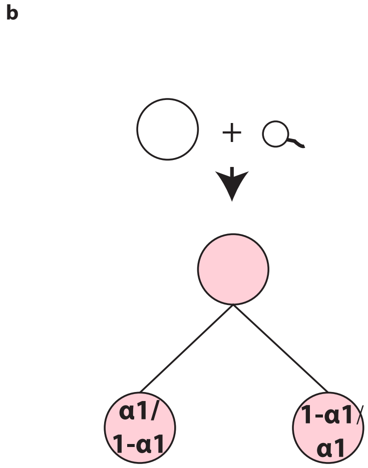


**VAF distribution supports an asymmetrical model of cell division. a,** Likelihood ratio-based 95% CI of the asymmetric cell contribution ratio is about 0.555:0.445-0.597:0.403 for the first cell generation. The 95% C.I. for $\alpha_1$ was calculated using the likelihood ratio (all values of $\alpha_1$ for which the likelihood drops off by no more than 1.92 units). **b,** A schematic diagram showing the asymmetric cell contribution.

**Supplementary Figure 11**

a

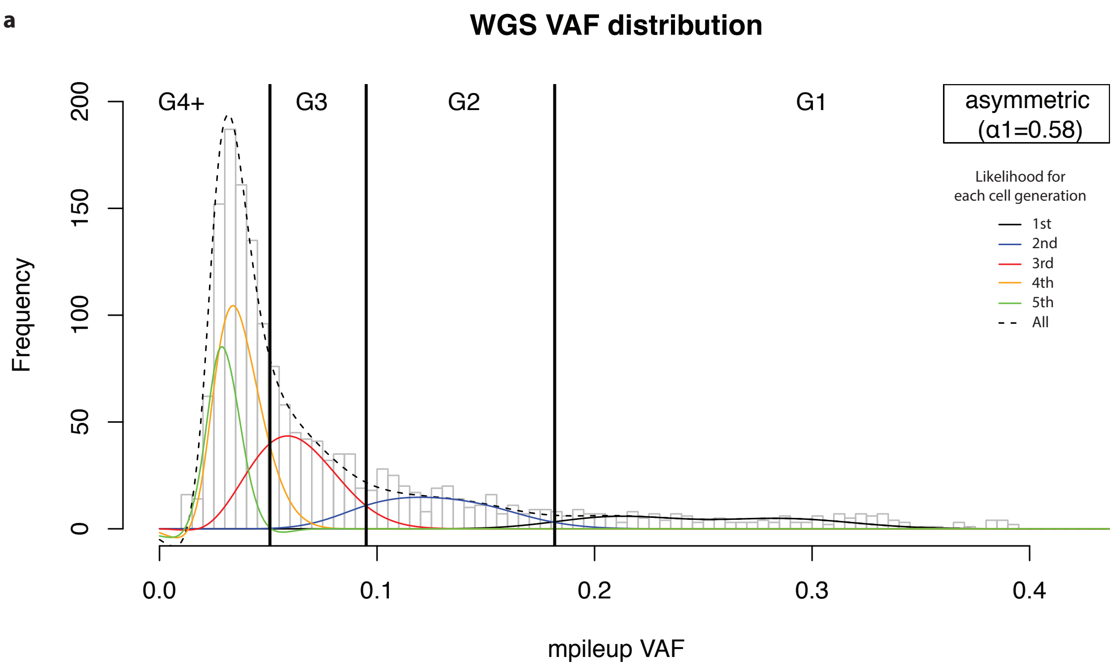


b

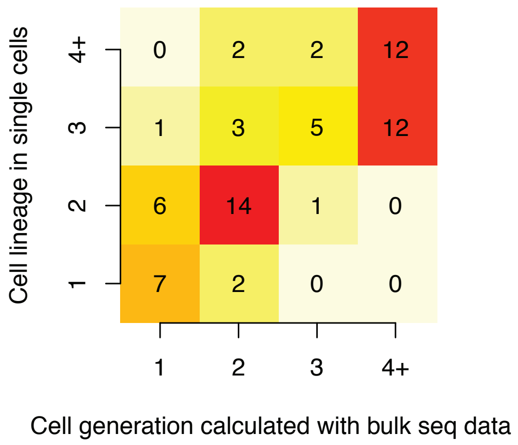


**Mutation assignments to cell generations under asymmetrical cell model. a,** Assignment of mosaic mutations to different cell generations in an asymmetrical cell model by a maximum likelihood approach. The likelihood curves for mosaics from different cell generations were estimated by considering mutation rate per cell generation, sensitivity curve of mosaic detection (for different real VAFs), frequencies of mosaic mutations with different VAFs after asymmetrical cell division as well as the binomial likelihood of an observed VAF under 250× read depth. See more details in the Methods. **b**, Cell generations for mosaics calculated with the asymmetric model were congruent with cell generations from cell lineages from single cell data in three individuals (UMB1465, UMB4638 and UMB4643).

#### Supplementary Figure 12

**a**

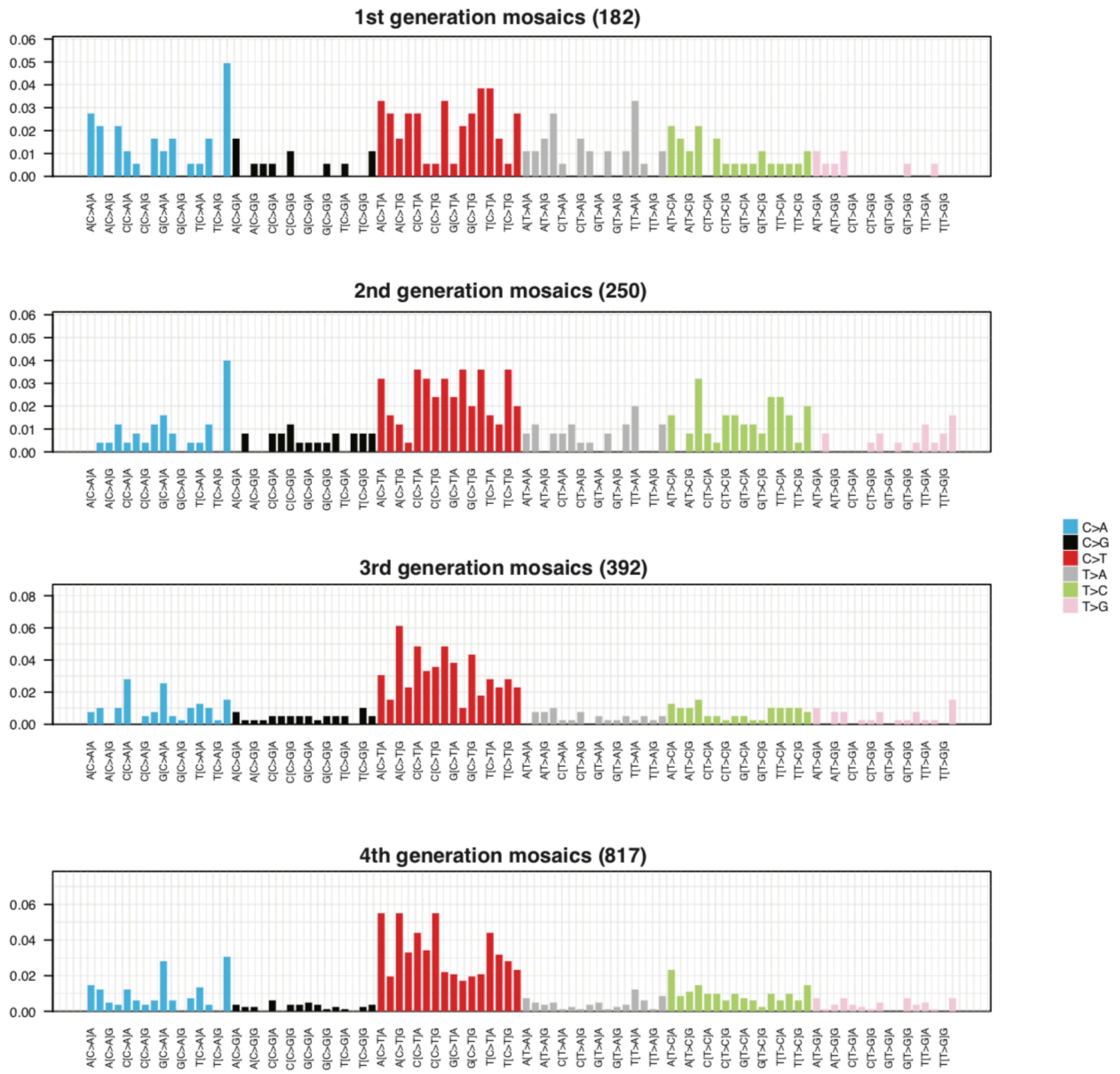


**b**

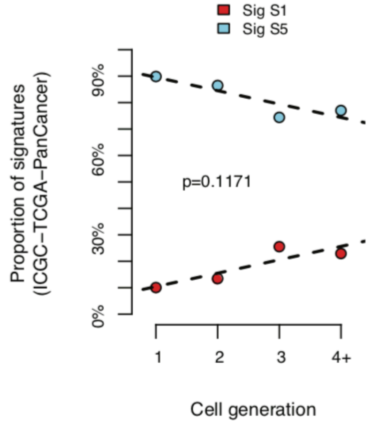


**Evolving spectra of somatic mutations in generations 1-4 of embryonic cell divisions. a,** C>T substitutions increase across cell generations while T>A substitutions decrease. **b,** PCAWG signature S1, which corresponds to COSMIC signature 1 and features CpG C>T mutations that increase in aging and cell division, trends toward increasing across cell divisions in our data.

#### Supplementary Figure 13

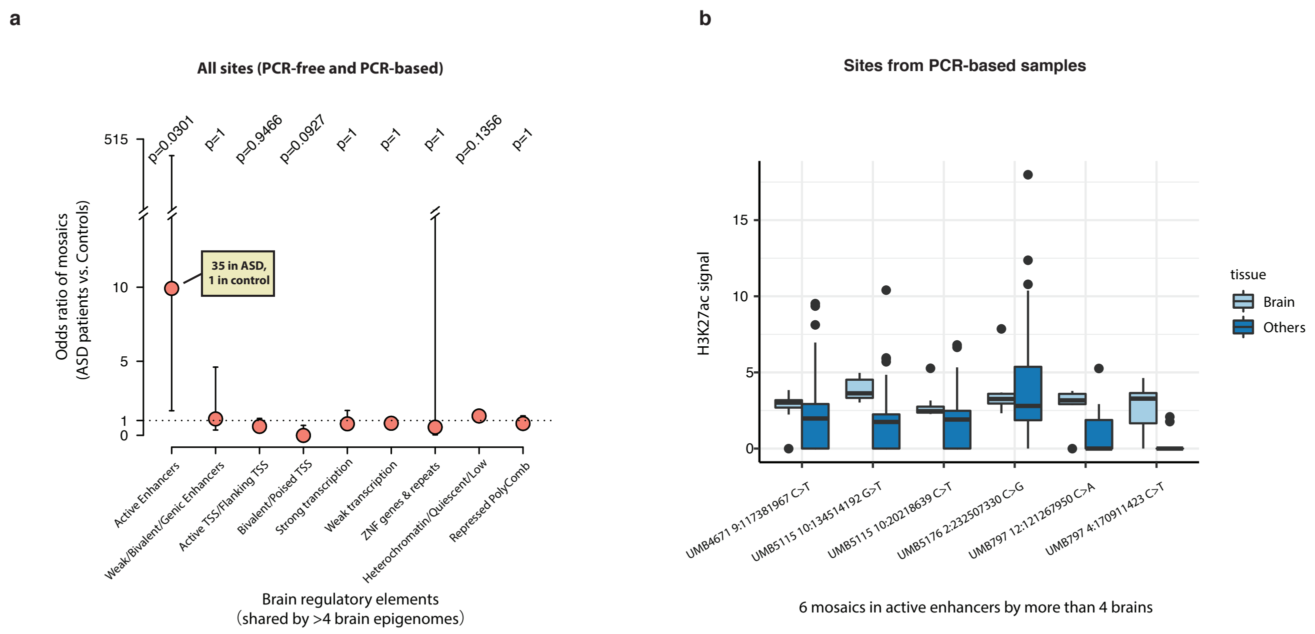


**ASD cases are enriched for mosaic mutations in brain-active enhancers when PCR-based mutations are included. a,** When both PCR-based and PCR-free mutations are included in the analysis, the excess of brain-active enhancer mutations in ASD cases remains significant. The odds ratio and error bars (95% CI) were calculated by Fisher's Exact test, and p-values were further corrected with Bonferroni correction. **b,** In total, 6 mutations from PCR-based samples were identified in enhancer regions that are brain-active in >50% (>4 of 8) brain epigenomes. The lower and upper hinges of the boxplot correspond to the first and third quartiles, and the middle lines correspond to the median values.

#### Supplementary Figure 14

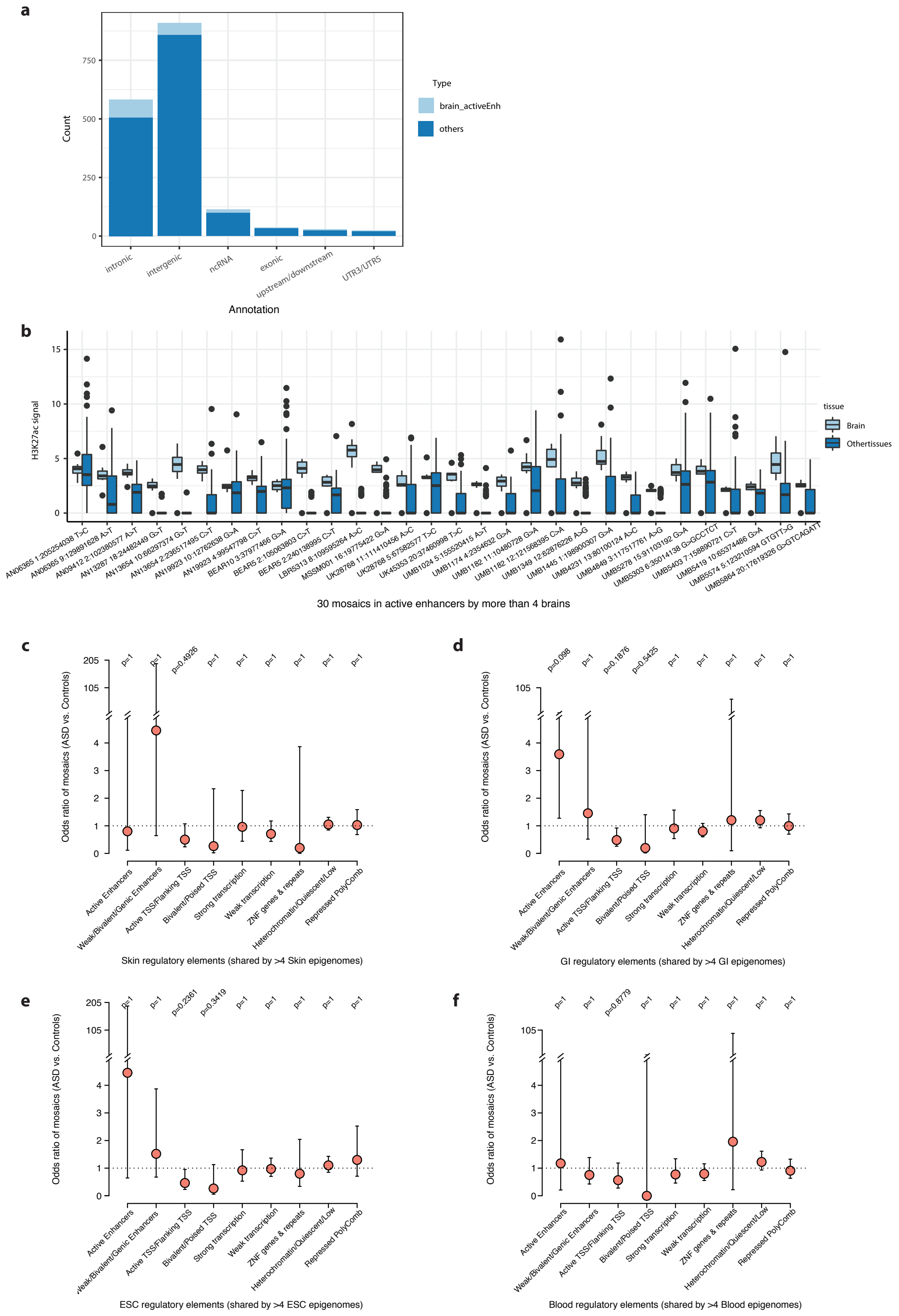


**ASD patients are not enriched for mosaic mutations in active enhancer regions in non-brain tissues. a**, Most of the brain-active enhancer mutations were within intronic and intergenic regions. **b**, Tissue-specific H3K27ac signals at the location of 30 mosaics (within brain active enhancer regions, shared by >4 Roadmap brain epigenomes, see Fig. 5a). A large fraction of the 30 mosaics have much higher H3K27ac signal in brain than other tissues, suggesting that these regions are brain-specific active enhancers. ASD cases are not enriched for mosaic mutations in any regulatory elements active in (**c**) skin, (**d**) GI tissues, (**e**) embryonic stem cells or (**f**) blood. All odds ratios and error bars (95% CI) in panels c-f were calculated by Fisher's Exact test, and p-values were further corrected with Bonferroni correction.

**Supplementary Figure 15**

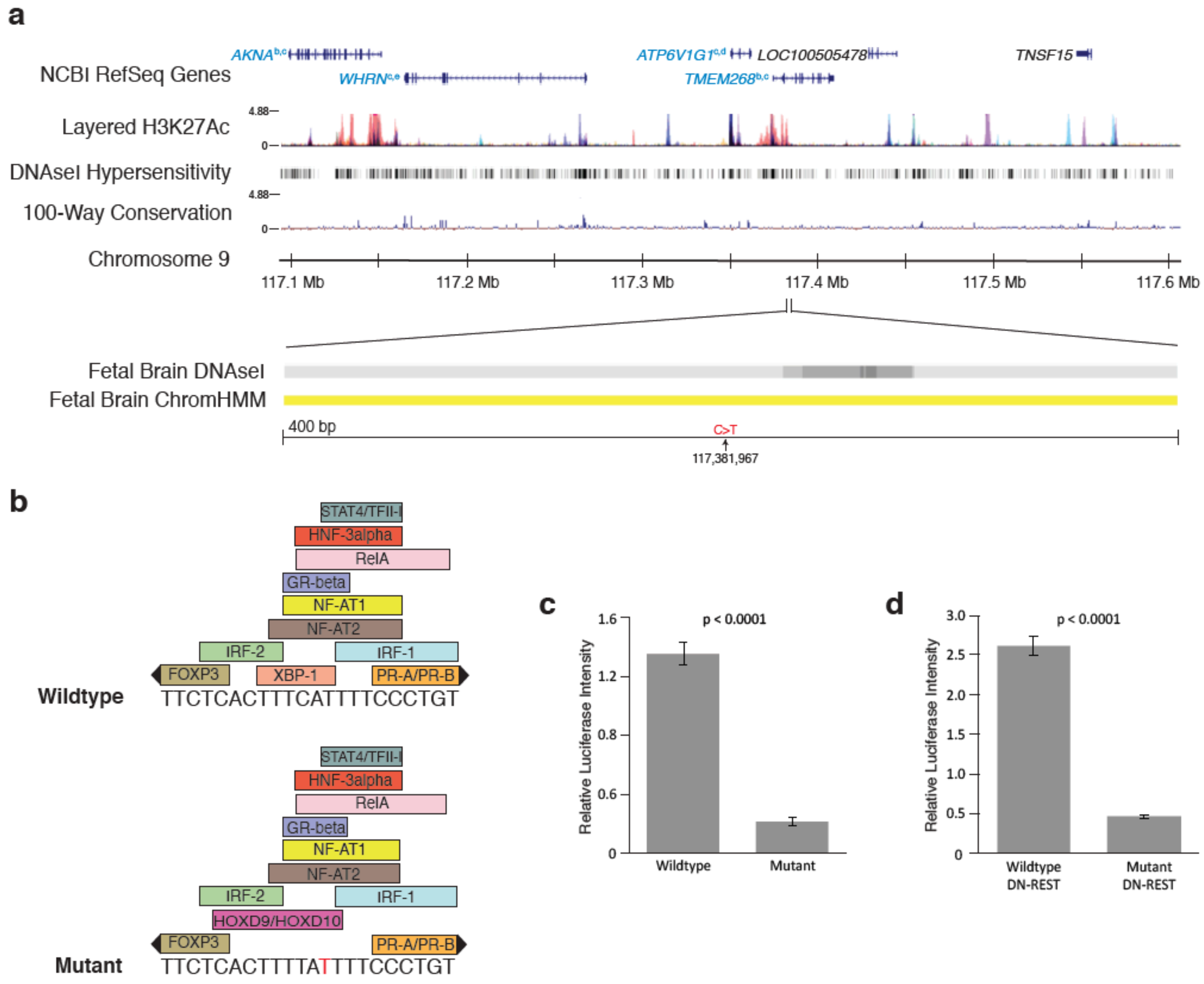


**Example of a mosaic mutation in ASD brain UMB4671 located in a brain-active enhancer. a,** Genes in blue type have functional evidence linking their expression to enhancer activity (a = Genotype Tissue Expression[44], b = Predicted Enhancer Targets[45], c = Hi-C sequencing data[46], d = Chromatin Interaction Analysis by Paired-End Tag Sequencing[47], e = Encode data). Yellow ChromHMM track represents weak active enhancer designation. **b,** The mutation is predicted to affect transcription factor binding. **c,** A mutant construct transfected into N2A cells results in reduced enhancer activity by 84% compared to wildtype construct (n=4, two-tailed t-test). **d,** In N2A cells pre-treated with DN-REST to assume a neuronal-like state, the mutant construct reduces enhancer activity by 82% (n=4, two-tailed t-test). Error bars represent 95% CIs.
